# The Drosophila *TART* transposon manipulates the piRNA pathway as a counter-defense strategy to limit host silencing

**DOI:** 10.1101/2020.02.20.957324

**Authors:** Christopher E. Ellison, Meenakshi S. Kagda, Weihuan Cao

## Abstract

Co-evolution between transposable elements (TEs) and their hosts can be antagonistic, where TEs evolve to avoid silencing and the host responds by reestablishing TE suppression, or mutualistic, where TEs are co-opted to benefit their host. The *TART-A* TE functions as an important component of Drosophila telomeres, but has also reportedly inserted into the *D. melanogaster* nuclear export factor gene *nxf2*. We find that, rather than inserting into *nxf2, TART-A* has actually captured a portion of *nxf2* sequence. We show that Nxf2 is involved in suppressing *TART-A* activity via the piRNA pathway and that *TART-A* produces abundant piRNAs, some of which are antisense to the *nxf2* transcript. We propose that capturing *nxf2* sequence allowed *TART-A* to target the *nxf2* gene for piRNA-mediated repression and that these two elements are engaged in antagonistic co-evolution despite the fact that *TART-A* is serving a critical role for its host genome.

## Introduction

Transposable elements (TEs) must replicate faster than their host to avoid extinction. The vast majority of new TE insertions derived from this replicative activity are deleterious to their host: they can disrupt and/or silence protein-coding genes and lead to chromosome rearrangements (Y. C. Lee, 2015; Y. C. G. Lee & Karpen, 2017; Petrov, Fiston-Lavier, Lipatov, Lenkov, & Gonzalez, 2011). In response to the mutational burden imposed by TEs, TE hosts have evolved elaborate genome surveillance mechanisms to identify and target TEs for suppression. One of the most well-known genome defense pathways in metazoan species involves the production of piwi-interacting small RNAs, also known as piRNAs (Brennecke et al., 2007). PiRNA precursors are produced from so-called piRNA clusters, which are located in heterochromatic regions of the genome and contain fragments of many families of TEs, whose insertions have accumulated in these regions. These precursors are processed into primary piRNAs, which use sequence homology to guide piwi-proteins to complementary transcripts produced by active transposable elements (Brennecke et al., 2007; Gunawardane et al., 2007). Piwi proteins induce transcriptional silencing through cleavage of the TE transcript. The sense-strand cleavage product of the TE transcript can then aid in processing piRNA precursors though a process known as the ping-pong cycle, which amplifies the silencing signal (Brennecke et al., 2007; Gunawardane et al., 2007). Alternatively, the cleaved transcript can be processed by the endonuclease Zucchini into additional “phased” piRNAs starting from the cleavage site and proceeding in the 3’ direction (Han, Wang, Li, Weng, & Zamore, 2015; Mohn, Handler, & Brennecke, 2015).

In addition to piRNAs, various other host mechanisms have evolved to target TEs (Cam, Noma, Ebina, Levin, & Grewal, 2008; Esnault et al., 2005; Satyaki et al., 2014; Thomas & Schneider, 2011)(mammalian systems reviewed in (Molaro & Malik, 2016)). Despite these multiple layers of genome surveillance, active TEs are found in the genomes of most organisms. The ubiquity of active TEs suggests that host silencing mechanisms are not completely effective, possibly because the TE and its host genome are involved in an evolutionary “arms race” where TEs are continuously evolving novel means to avoid host silencing and the host genome is constantly reestablishing TE suppression (Parhad & Theurkauf, 2019). On the host side, many TE silencing components have been shown to be evolving rapidly under positive selection (Crysnanto & Obbard, 2019; Helleu & Levine, 2018; Jacobs et al., 2014; Kelleher, Edelman, & Barbash, 2012; Kolaczkowski, Hupalo, & Kern, 2011; Levine, Vander Wende, Hsieh, Baker, & Malik, 2016; Obbard, Jiggins, Bradshaw, & Little, 2011; Obbard, Jiggins, Halligan, & Little, 2006; Simkin, Wong, Poh, Theurkauf, & Jensen, 2013), in agreement with on-going host-TE conflict. On the transposon side, a TE can mount a counter-defense by silencing or blocking host factors (Fu et al., 2013; McCue, Nuthikattu, & Slotkin, 2013; Nosaka et al., 2012) or simply evade host silencing by replicating in permissive cells (L. Wang, Dou, Moon, Tan, & Zhang, 2018) or cloaking themselves in virus-like particles (Mari-Ordonez et al., 2013). However, there are surprisingly few examples of any of these strategies (Cosby, Chang, & Feschotte, 2019). In fact, there is some evidence that, rather than an evolutionary arms race, the rapid evolution of host silencing genes is related to avoiding gene silencing due to off-target effects (i.e. piRNA autoimmunity (Blumenstiel, Erwin, & Hemmer, 2016; Luyang Wang, Barbash, & Kelleher, 2019)) and/or co-evolution with viruses (reviewed in (Cosby et al., 2019)).

While there are currently only a few examples of TE counter-defense strategies, there are many examples of TEs being co-opted by their host genome for its own advantage (see reviews (Bohne, Brunet, Galiana-Arnoux, Schultheis, & Volff, 2008; Chuong, Elde, & Feschotte, 2017; Cosby et al., 2019; Feschotte, 2008; Volff, 2006)). TEs can disperse regulatory sequences across the genome, which allows them to rewire gene regulatory networks. Such rewiring phenomena have been implicated in a variety of evolutionary innovations from pregnancy to dosage compensation (Chuong, Elde, & Feschotte, 2016; Chuong, Rumi, Soares, & Baker, 2013; Dunn-Fletcher et al., 2018; C. Ellison & Bachtrog, 2019; C. E. Ellison & Bachtrog, 2013; Fuentes, Swigut, & Wysocka, 2018; Lynch, Leclerc, May, & Wagner, 2011; Lynch et al., 2015; Notwell, Chung, Heavner, & Bejerano, 2015; Pontis et al., 2019). TEs are also an important source of host genes and noncoding RNAs (Joly-Lopez & Bureau, 2018; Kapusta et al., 2013). Hundreds of genes in species ranging from mammals to plants have been acquired from transposons (Bohne et al., 2008; Joly-Lopez, Hoen, Blanchette, & Bureau, 2016; Volff, 2006). Finally, TEs can act as structural components of the genome. There is evidence that TEs may play a role in centromere specification in a variety of species (Chang et al., 2019; Chueh, Northrop, Brettingham-Moore, Choo, & Wong, 2009; Klein & O’Neill, 2018), and in Drosophila, which lacks telomerase, specific TEs serve as telomeres by replicating to chromosome ends (Levis, Ganesan, Houtchens, Tolar, & Sheen, 1993; Traverse & Pardue, 1988).

In *Drosophila melanogaster*, three related non-LTR retrotransposons occupy the telomeres: *HeT-A, TAHRE*, and *TART*, which are often abbreviated as HTT elements (Abad et al., 2004b; Biessmann et al., 1992; Levis et al., 1993; Sheen & Levis, 1994). These elements belong to the Jockey clade of Long Interspersed Nuclear Elements (LINEs), which contain open reading frames for gag (ORF1) and an endonuclease/reverse transcriptase protein (ORF2, lost in *HeT-A*) (Malik, Burke, & Eickbush, 1999; Villasante et al., 2007). These elements form head-to-tail arrays at the chromosome ends and their replication solves the chromosome “end-shortening” problem without the need for telomerase (Biessmann & Mason, 1997).

These telomeric elements represent a unique case of TE domestication. They serve a critical role for their host genome, yet they are still active elements, capable of causing mutational damage if their activity is left unchecked (Khurana, Xu, Weng, & Theurkauf, 2010; Savitsky, Kravchuk, Melnikova, & Georgiev, 2002; Savitsky, Kwon, Georgiev, Kalmykova, & Gvozdev, 2006). All three elements have been shown to produce abundant piRNAs, and RNAi knockdown of piRNA pathway components leads to their upregulation (Savitsky et al., 2006; Shpiz & Kalmykova, 2011; Shpiz et al., 2011), consistent with the host genome acting to constrain their activity and raising the possibility that, despite being domesticated, these elements are still in conflict with their host (Y. C. Lee, Leek, & Levine, 2017).

There are multiple lines of evidence that this is indeed the case: the protein components of Drosophila telomeres are rapidly evolving under positive selection, potentially due to a role in preventing the HTT elements from overproliferation (Y. C. Lee et al., 2017). There is a high rate of gain and loss of HTT lineages within the melanogaster species group (Saint-Leandre, Nguyen, & Levine, 2019), and there is dramatic variation in telomere length among strains from the Drosophila Genetic Reference Panel (DGRP) (Wei et al., 2017). These observations are more consistent with evolution under conflict rather than a stable symbiosis (Saint-Leandre et al., 2019). Furthermore, the nucleotide sequence of the HTT elements evolves extremely rapidly, especially in their unusually long 3’ UTRs (Casacuberta & Pardue, 2002; Danilevskaya, Tan, Wong, Alibhai, & Pardue, 1998). Within *D. melanogaster*, three *TART* subfamilies have been identified which contain completely different 3’ UTRs, and which are known as *TART-A, TART-B*, and *TART-C* (Sheen & Levis, 1994).

In this study we have characterized the presence of sequence within the coding region of the *D. melanogaster nxf2* gene that was previously annotated as an insertion of the *TART-A* transposon (Sackton et al., 2009). We find that the shared homology between *TART-A* and *nxf2* is actually the result of *TART-A* acquiring a portion of the *nxf2* gene, rather than the *nxf2* gene gaining a *TART-A* insertion. We also find that *nxf2* plays a role in suppressing *TART-A* activity, likely via the piRNA pathway. Our findings support a model where *TART-A* produces antisense piRNAs that target *nxf2* for suppression as a counter-defense strategy in response to host silencing. We identified *nxf2* cleavage products from degradome-seq data that are consistent with Aub-directed cleavage of *nxf2* transcripts and we find that, across the Drosophila Genetic Reference Panel (DGRP), *TART-A* copy number is negatively correlated with *nxf2* expression. Our findings suggest that TEs can selfishly manipulate host silencing pathways in order to increase their own copy number and that a single TE family can benefit, as well as antagonize, its host genome.

## Results

### The *TART*-like region of *nxf2* is conserved across the melanogaster group

It was previously reported that the homology between *nxf2* and *TART-A* is due to an insertion of the *TART-A* transposable element in the *nxf2* gene that became fixed in the ancestor of *D. melanogaster* and *D. simulans* (Sackton et al., 2009). To investigate the homology between these elements in more detail, we first extracted 700 bp of sequence from the 3’ region of the *nxf2* gene that was annotated as a *TART-A* insertion (**Figure 1A**) and used BLAST (Altschul, Gish, Miller, Myers, & Lipman, 1990) to search this sequence against the *TART-A* RepBase sequence, which was derived from a full-length *TART-A* element cloned from the *iso1 D. melanogaster* reference strain (Abad et al., 2004a). Within the 700 bp segment of *nxf2*, there are four regions of homology between it and the 3’ UTR of the *TART-A* consensus sequence. These regions are between 63 bp and 228 bp in length and 93% - 96% sequence identity (**Figure 1B**). The 5’ UTR of *TART-A* is copied from its 3’ UTR during reverse transcription, which means that, for a given element, both UTRs are identical in sequence (George, Traverse, DeBaryshe, Kelley, & Pardue, 2010). The homology with the *nxf2* 3’ UTR is therefore mirrored in the 5’ UTR as well (**Figure 1B**).

**Figure 1.**
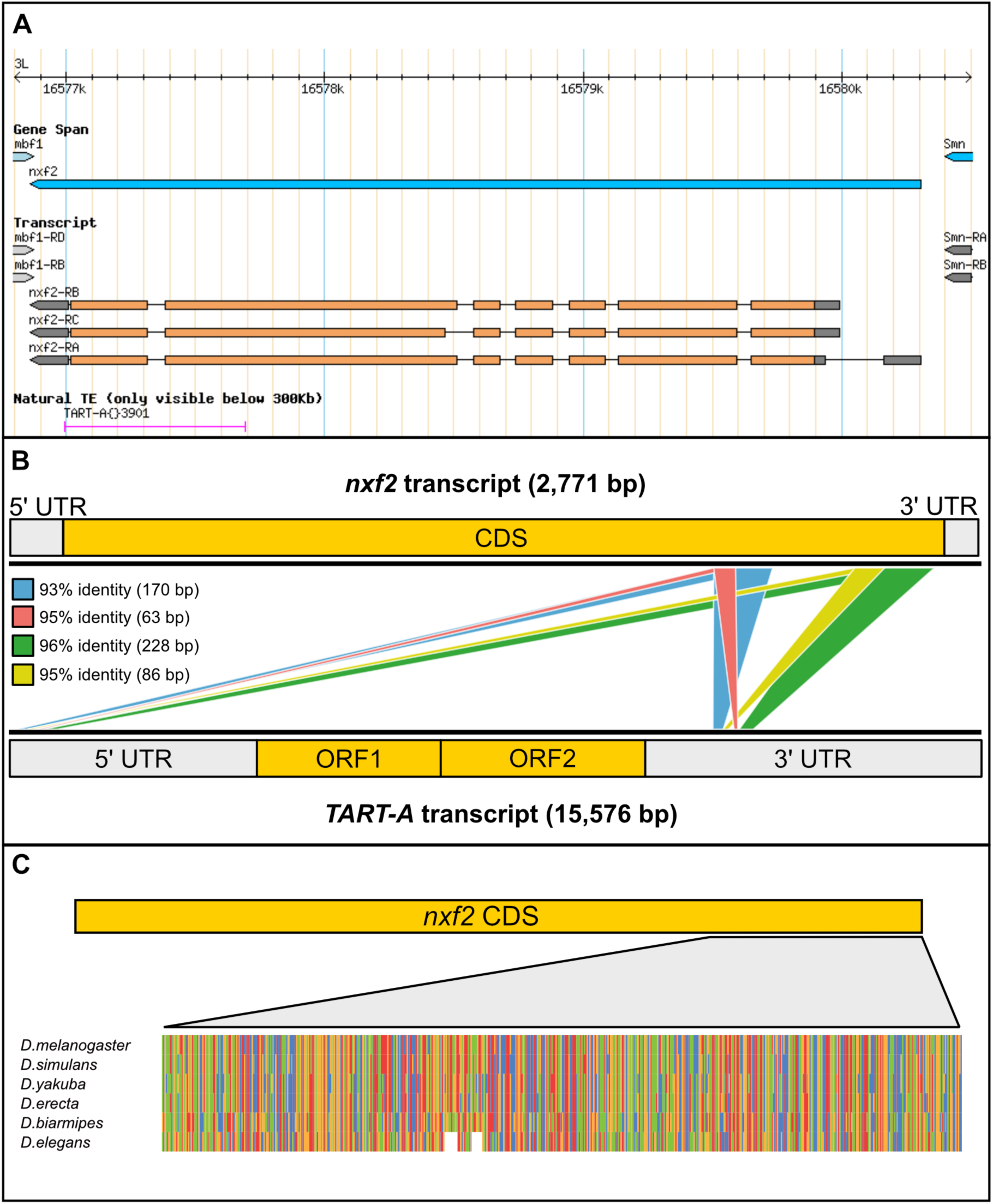
Shared homology between the *D. melanogaster nxf2* gene and the *TART-A* transposable element. (A) GBrowse screenshot from FlyBase showing the *nxf2* gene model along with the annotated *TART-A* TE insertion. Note that the *TART-A* annotation overlaps the 3’ coding sequence of *nxf2*. (B) BLAST hits between the RepBase *TART-A* sequence and the *nxf2* transcript. Each colored box represents a single BLAST alignment. The 5’ UTR of *TART-A* is copied from its 3’ UTR during replication. The two UTRs are therefore identical in sequence and the homology between *nxf2* and the *TART-A* 3’ UTR is mirrored in the 5’ UTR. (C). A zoomed-out multiple sequence alignment of *nxf2* orthologs for six species from the melanogaster species group shows that the *TART-like* region of *nxf2* is present in all six species.

To investigate the evolutionary origin of the homology between *nxf2* and *TART-A*, we identified *nxf2* orthologs in *D. simulans, D. yakuba, D. erecta, D. biarmipes*, and *D. elegans*. We created a multiple sequence alignment and extracted the sub-alignment corresponding to the 700 bp segment with homology to *TART-A* (**Figure 1C**). The *TART*-like region of *nxf2* is clearly present in all six of these species, which means that, if this portion of the *nxf2* gene was derived from an insertion of a *TART-A* element, the most recent timepoint at which the insertion could have occurred is in the common ancestor of the melanogaster group, ∼15 million years ago (Obbard et al., 2012). At the nucleotide level, there is only weak homology between *nxf2* coding sequence and transcripts from more distantly related Drosophila species, such as *D. pseudoobscura*. However, at the peptide level, the C-terminal region of Nxf2, which was thought to be derived from *TART-A*, is actually conserved across Drosophila, from *D. melanogaster* to *D. virilis* (**Figure S1**), suggesting that, if a *TART-A* element did insert into the *nxf2* gene, it was not a recent event.

### A portion of *nxf2* was captured by the *D. melanogaster TART-A* element

If an ancestral *TART-A* element was inserted into the *nxf2* gene in the common ancestor of the melanogaster group, the shared homology between *nxf2* and *TART-A* should be present in most, if not all, extant species in the group. To test this prediction, we obtained the sequences for previously identified *TART-A* homologs from *D. yakuba* and *D. sechellia* (Casacuberta & Pardue, 2002; Villasante et al., 2007). We aligned these sequences to the *D. melanogaster TART-A* consensus sequence and found that the *TART-A* region that shares homology with the *nxf2* gene is only present in the *D. melanogaster TART-A* sequence (**Figure 2A & S2**). Next, we used BLAST to search the canonical *TART-A* sequence against the *D. melanogaster* reference genome. We identified 5 full-length *TART-A* sequences in the assembly (3 from the X chromosome and 2 from the dot chromosome), all of which contain the *nxf2*-like sequence. The *nxf2*-like sequence from these five elements is 100% identical to that from the canonical *TART-A* sequence. We also identified an additional four *TART-A* fragments that overlapped with the *nxf2*-like region. One of the four is also 100% identical to the canonical sequence while the remaining three are between 96%-99% identical to the canonical sequence.

**Figure 2.**
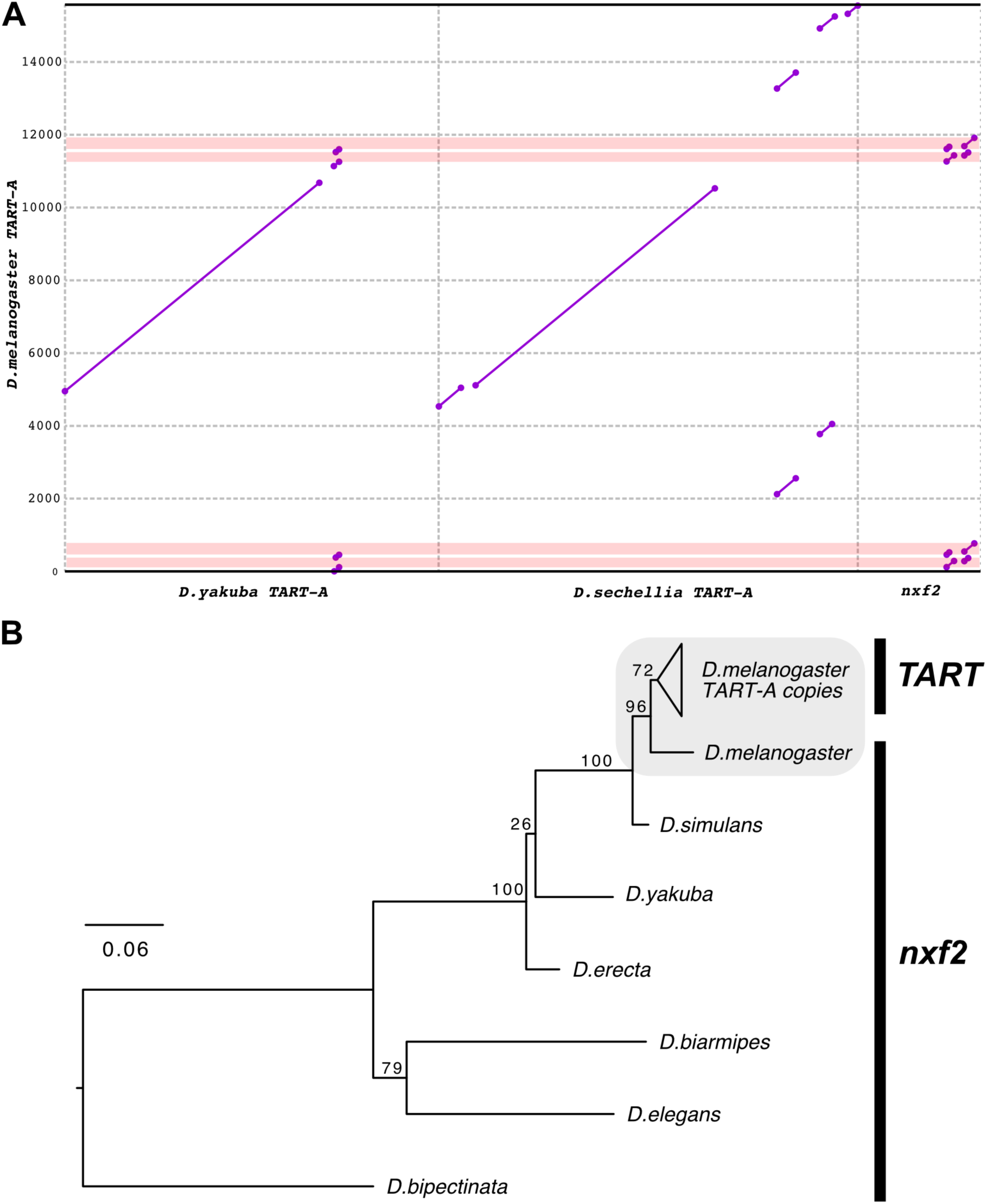
The *TART-A*/*nxf2* homology is unique to *D. melanogaster*. (A) Dotplot comparing *D. melanogaster TART-A* to its homologs in *D. yakuba* and *D. sechellia*. The diagonal lines denote regions of homology while the pink boxes show the location of the *nxf2*-like sequence in the *D. melanogaster TART-A*. Neither the *D. yakuba* nor the *D. sechellia TART-A* sequences contain *nxf2*-like sequence. However, the regions directly flanking the *nxf2*-like sequence in *D. melanogaster* are also present in *D. yakuba* (see **Figure S2** for magnified view). (B) Gene tree showing relative age of shared homology. We aligned the *nxf2*-like sequences from nine copies of *TART-A* in the *D. melanogaster* reference genome to the *nxf2* transcripts from six Drosophila species and inferred a maximum likelihood phylogeny using RAxML. *D. melanogaster nxf2* is most closely related to the *nxf2*-like sequences present in the *D. melanogaster TART-A* copies, suggesting the shared homology occurred after the divergence between *D. melanogaster* and *D. simulans*.

We added these nine sequences to the multiple sequence alignment in **Figure 1C** and inferred a maximum likelihood phylogeny in order to better understand the evolutionary history of the *nxf2*/*TART* shared homology (**Figure 2B**). The youngest node in the phylogeny represents the split between the *D. melanogaster nxf2* and *TART-A* elements, suggesting that the event leading to the shared homology between these sequences occurred relatively recently, which is consistent with the high degree of sequence similarity between the *D. melanogaster TART-A* and *nxf2* subsequences. Based on these results, we conclude that the *nxf2/TART-A* shared homology is much more likely to have arisen via the recent acquisition of *nxf2* sequence by *TART-A* after the split of *D. melanogaster* from *D. simulans*/*sechellia*, rather than an insertion of *TART-A* into the *nxf2* gene. The mechanism by which *TART-A* could have acquired a portion of *nxf2* is not clear, however one possibility is via transduction, a process where genomic regions flanking a TE insertion can be incorporated into the TE itself due to aberrant retrotransposition (Moran, DeBerardinis, & Kazazian, 1999; Pickeral, Makalowski, Boguski, & Boeke, 2000).

### The *nxf2* gene plays a role in suppressing the activity of *D. melanogaster* telomeric elements

Nxf2 is part of an evolutionarily conserved gene family with functions related to export of RNA from the nucleus (Herold et al., 2000). In Drosophila, a paralog of *nxf2* (*nxf1*) has been shown to be involved in the nuclear export of piRNA precursors and the *nxf2* gene itself was identified as a member of the germline piRNA pathway via an RNAi screen (Czech, Preall, McGinn, & Hannon, 2013; Dennis, Brasset, Sarkar, & Vaury, 2016). More recently, several studies have independently shown that Nxf2 is involved in the co-transcriptional silencing of transposons as part of a complex with Nxt1 and Panoramix (Batki et al., 2019; Fabry et al., 2019; Murano et al., 2019; Zhao et al., 2019). To determine whether *nxf2* is involved in the suppression of *TART-A*, we used a short hairpin RNA (shRNA) from the Drosophila transgenic RNAi project (TRiP) with a nos-GAL4 driver to target and knockdown expression of *nxf2* in the ovaries. We sequenced total RNA from the *nxf2* knockdown and a control knockdown of the *white* gene. We observed a strong increase in expression for a variety of TE families upon knockdown of *nxf2* (**Figure S3**). The three telomeric elements *HeT-A, TAHRE*, and *TART-A*, are among the top 10 most highly upregulated transposable elements, with *HeT-A* showing ∼300-fold increase in expression in the *nxf2* knockdown (*TAHRE*: ∼110-fold increase, *TART-A*: ∼30-fold increase)(**Figure 3**). We repeated the experiment using a shRNA that targeted a different region of *nxf2* and observed a similar pattern and strong correlation between TE expression profiles of both knockdowns (Spearman’s rho=0.94, **Figure S4**). These results support previous findings that *nxf2* is a component of the germline piRNA pathway and show that this gene is particularly important for the suppression of the telomeric TEs *HeT-A, TAHRE*, and *TART-A*.

**Figure 3.**
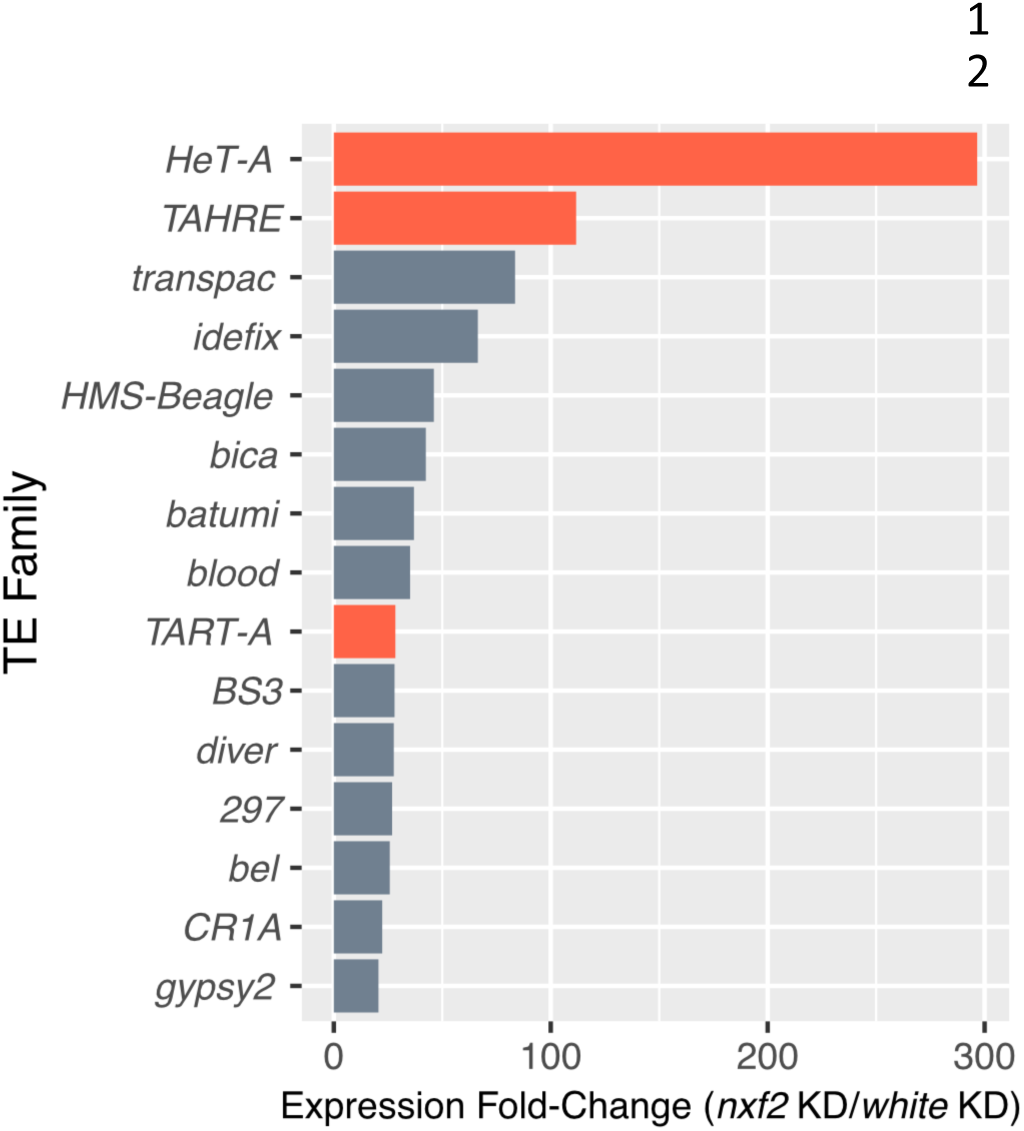
RNAi knockdown of *nxf2* leads to strong upregulation of HTT elements. We examined TE expression profiles using RNA-seq of total RNA from ovaries in a *nxf2* knockdown versus a control knockdown of the *white* gene. We found that a variety of transposable elements show increased expression in the *nxf2* knockdown (see **Figure S3** for all TEs), however the three telomeric HTT elements (red bars) are among the top 10 most highly upregulated TEs.

### *TART-A* piRNAs may target *nxf2* for silencing

Previous studies have reported abundant piRNAs derived from the telomeric TEs, *HeT-A, TAHRE* and *TART-A* (Savitsky et al., 2006; Shpiz et al., 2007; Shpiz et al., 2011). We sought to determine whether piRNAs arising from the *nxf2*-like region of *TART-A* could be targeting the *nxf2* gene for downregulation via the piRNA pathway. We used previously published piRNA data from 16 wild-derived strains from the Drosophila Genetic Reference Panel (DGRP)(Song et al., 2014). Because the 5’ UTR is copied from the 3’ UTR, we masked the 5’ UTR of *TART-A* before aligning the piRNA data. Among the 16 strains, we found a large variation in *TART-A* piRNA production ranging from 60 – 12,300 reads per million (RPM). From the pool of 16 strains, we identified ∼1.3 million reads that aligned to *TART-A*, 98% of which map uniquely (see Methods)(**Figure 4A**). *TART-A* piRNAs have previously been shown to exhibit the 10bp overlap signature of ping-pong cycle amplification (Hur et al., 2016) and we identified both sense and antisense piRNAs arising from *TART-A* (**Figure 4B**) as well as an enrichment of alignments where the 5’ end of one piRNA is found directly after the 3’ end of the previous piRNA (i.e. 3’ to 5’ distance of 1), consistent with piRNA phasing (**Figure 4C**). We identified ∼95,000 piRNAs arising from the *TART-A* region that shares homology with *nxf2*. Of these reads, 59% are antisense to *TART-A* and 41% are sense.

**Figure 4.**
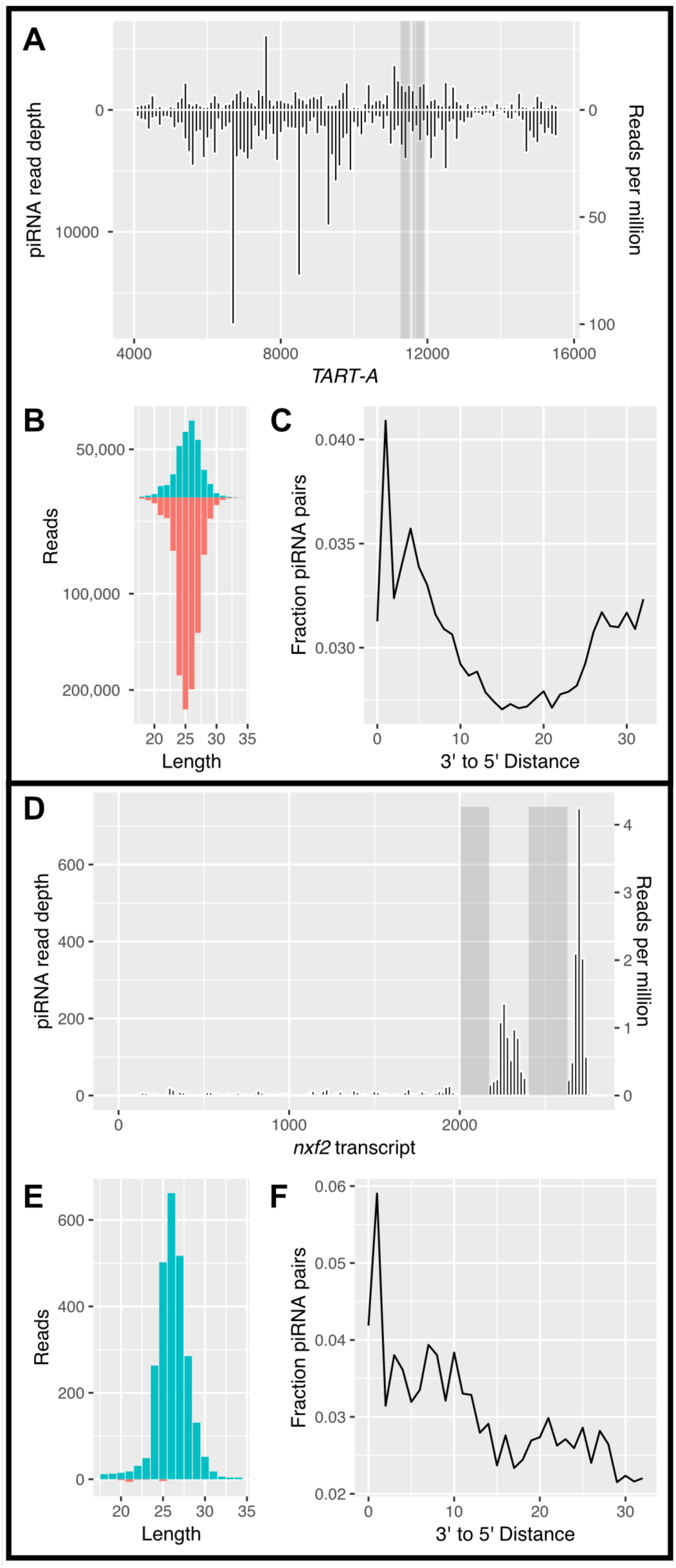
piRNAs are produced from both *TART-A* and *nxf2*. (A) We aligned previously published piRNA data from the *D. melanogaster* Drosophila Genetic Reference Panel (DGRP)(Song et al., 2014) to *TART-A* and examined read coverage across the element. We find abundant sense and antisense piRNA production across most of the element, including the regions containing the *nxf2*-like sequence (grey boxes). Note that the 5’ UTR of *TART-A* is copied from the 3’ UTR during replication and is therefore identical in sequence. We masked the 5’ UTR (positions 1-4000) for this analysis. (B) The length of aligned reads are consistent with that expected for piRNAs and the *TART-A* derived piRNAs are biased towards the minus strand. (C) *TART-A* piRNAs show an enrichment of alignments where the 5’ end of one piRNA is found directly after the 3’ end of the previous piRNA (i.e. distance of 1), consistent with piRNA phasing. (D) Unlike *TART-A, nxf2* produces piRNAs primarily in the regions directly downstream from its *TART*-like sequence (grey boxes). The vast majority of these piRNAs are only from the sense strand of *nxf2* (panel E) and also show the signature of phasing (panel F). Note that the *TART*-like sequence of *nxf2* was masked for this analysis to avoid cross-mapping of *TART*-derived piRNAs to the *nxf2* transcript.

We next focused on piRNA production from *nxf2*. We reasoned that, if *nxf2* expression is subject to piRNA-mediated regulation, we should see piRNAs derived from the *nxf2* transcript, outside of the region that shares homology with *TART-A*. We masked the *nxf2*/*TART-A* region of shared homology and aligned the piRNA sequence data to the *nxf2* transcript. We found low but consistent production of piRNAs from *nxf2* across all 16 DGRP strains (between 1.5 and 41 RPM), with 99.7% of *nxf2*-aligned reads mapping uniquely. To increase sequencing depth, we pooled the data from all 16 strains (2,624 *nxf2* reads total) and examined piRNA abundance along the *nxf2* transcript (**Figure 4D**). We found that the most abundant production of piRNAs from *nxf2* occurs at the 3’ end of the transcript, downstream from the regions of shared homology with *TART-A* (**Figure 4D**). Overall, 99.4% of reads from *nxf2* are derived from the sense strand of the transcript (**Figure 4E**) and the *nxf2* piRNAs also show evidence of phasing (**Figure 4F**). The enrichment of *nxf2*-derived piRNAs downstream from the region of shared homology with *TART-A*, along with our observation that almost all *nxf2* piRNAs are derived from the sense strand, suggests that these piRNAs are not amplified via the ping-pong cycle, but are instead produced by the Zucchini-mediated phasing process.

These results are consistent with a model where antisense piRNAs from the *nxf2*-like region of *TART-A* are bound by Aubergine and targeted to sense transcripts from the *nxf2* gene. Aub cleaves target transcripts between the bases paired to the 10^th^ and 11^th^ nucleotides of its guide piRNA, resulting in a cleavage product with a 5’ monophosphate that shares a 10 bp sense:antisense overlap with the guide piRNA that triggered the cleavage. These cleavage products can be enriched and sequenced using an approach known as degradome-seq (Addo-Quaye, Eshoo, Bartel, & Axtell, 2008). We analyzed published degradome-seq and Aub-immunoprecipitated piRNA data from wild-type *D. melanogaster* ovaries (W. Wang et al., 2014) to determine whether we could detect *nxf2* cleavage products resulting from targeting by antisense *TART-A* piRNAs. The degradome-seq data are 100 bp paired-end reads which are long enough to distinguish between the *TART*-like region of *nxf2* and the *nxf2*-like region of *TART-A*. We found three locations within the *TART*-like region of *nxf2* where we observe degradome cleavage products that share the characteristic 10bp sense:antisense overlap with *TART-A* antisense piRNAs (**Figure S5**). These results can be explained under the following model: *TART-A* antisense piRNAs are produced by the ping-pong cycle and bound to Aubergine. A subset of these piRNAs (those from the *nxf2*-like region of *TART-A*) guide Aub to *nxf2* transcripts which are then cleaved. Aub cleavage products can be further processed by Zucchini in the 5’ to 3’ direction thereby producing phased piRNAs from *nxf2* transcripts downstream from the *nxf2*/*TART-A* regions of shared homology (**Figure 5**).

**Figure 5.**
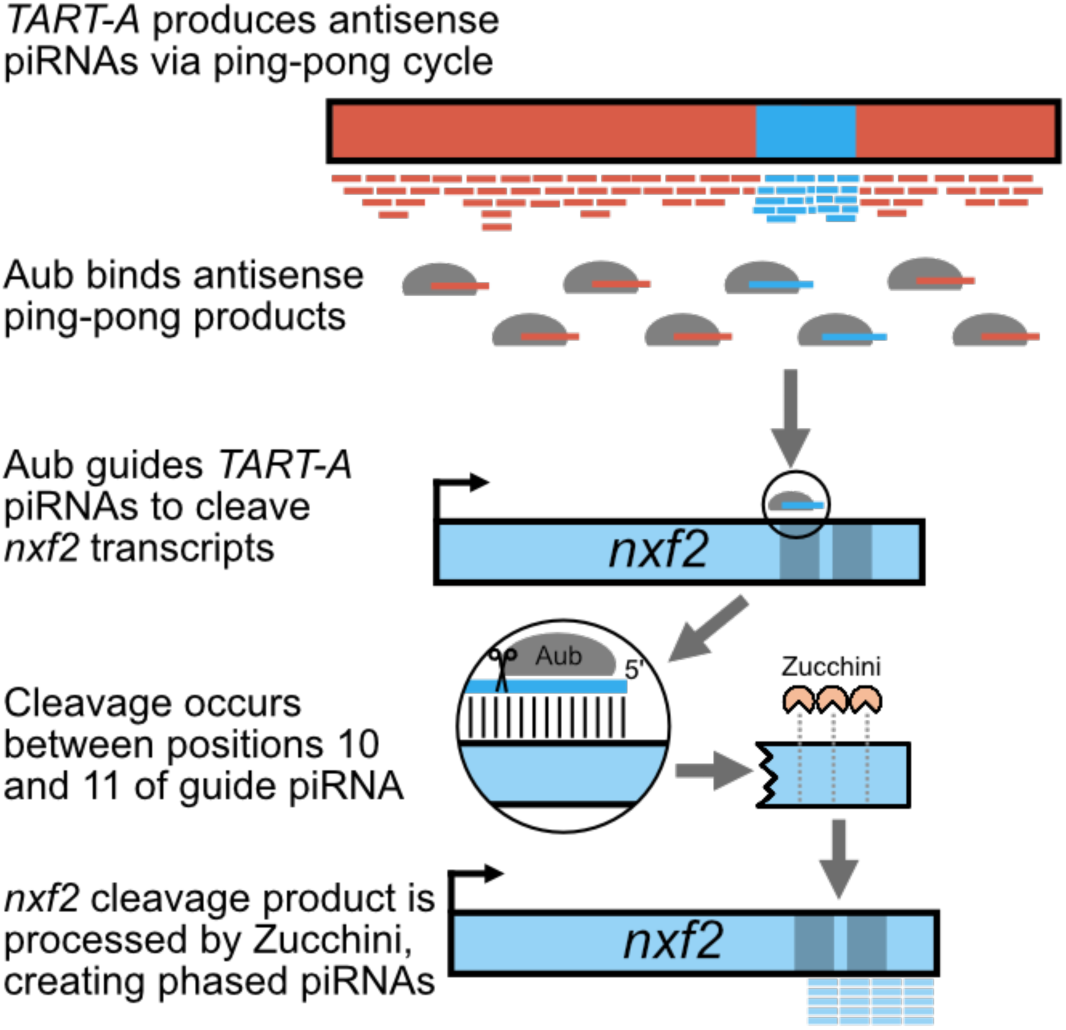
Model describing generation of phased piRNAs from *nxf2*. *TART-A* produces abundant antisense piRNAs derived from ping-pong amplification, including from the *TART-A*/*nxf2* region of shared homology (blue box on red background). The PIWI protein Aubergine binds antisense ping-pong piRNAs, a subset of which share homology with *nxf2*. These piRNAs guide Aub to *nxf2* and result in cleavage of the transcript between the 10^th^ and 11^th^ nucleotide of the guide piRNA. Transcript cleavage creates an *nxf2* cleavage product that shares a 10 bp sense:antisense overlap with the guide piRNA (see **Figure S5**). The *nxf2* cleavage product can by subsequently processed by the Zucchini endonuclease, creating phased piRNAs starting from the site of Aub cleavage and proceeding to the 3’ end of the *nxf2* transcript.

If piRNAs from *TART-A* are targeting *nxf2* and downregulating its expression, knockdown of piRNA pathway components that either decrease piRNA production from *TART-A* (ping-pong and/or primary piRNA pathway components) or disrupt silencing of *nxf2* (primary piRNA components) should result in an increase in expression of *nxf2*. We analyzed published RNA-seq data from nos-GAL4 driven knockdowns of sixteen genes that were identified as components of the piRNA pathway and that were specifically shown to be involved in repression of *HeT-A* and *TAHRE* (Czech et al., 2013). We compared the expression of *nxf2* in each piRNA component knockdown to its expression in the control knockdown of the *white* gene and found that *nxf2* shows increased expression in 14 of the 16 knockdowns, which represents a significant skew towards upregulation (one-sided binomial test P=0.002)(**Figure 6**).

**Figure 6.**
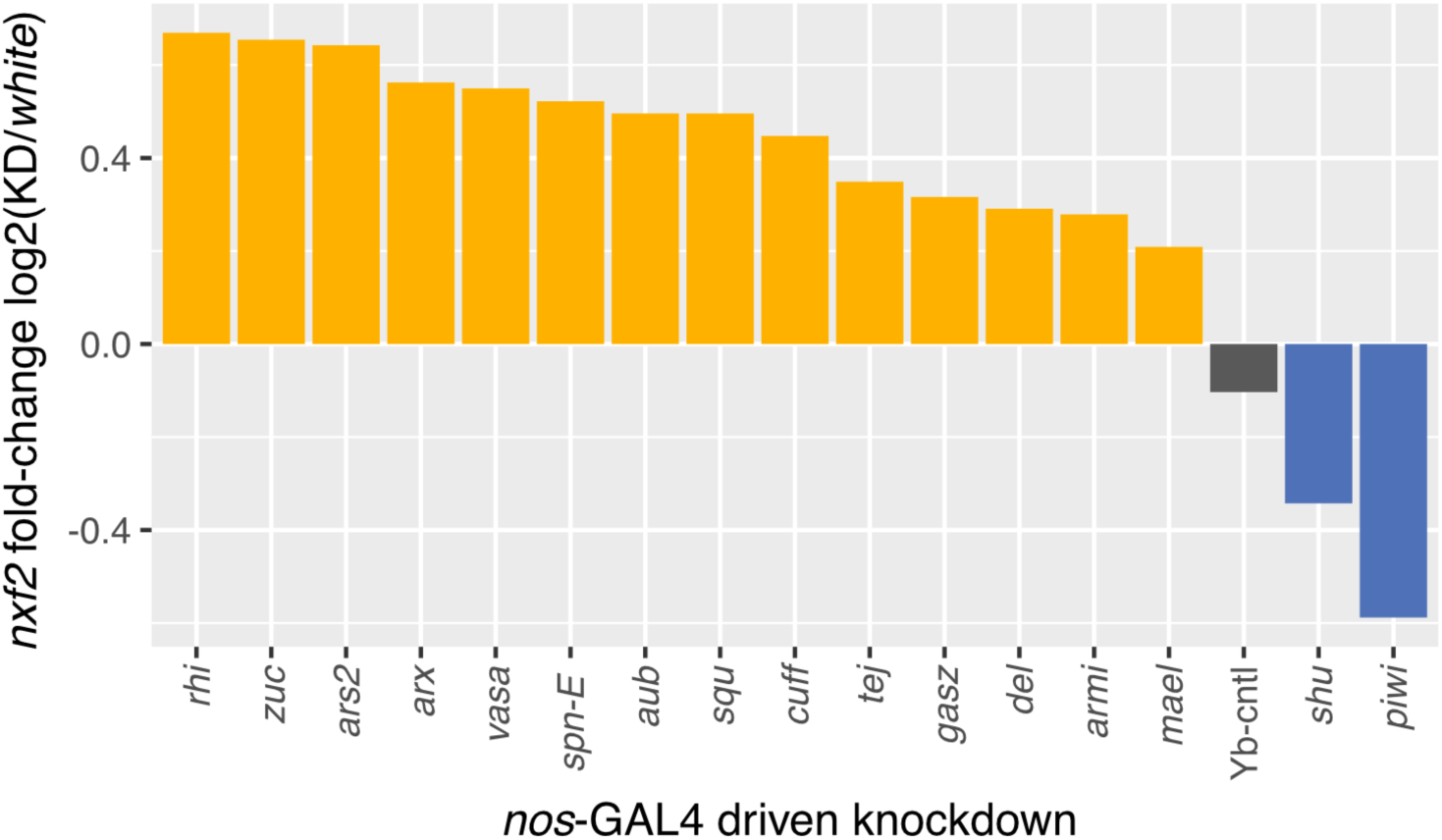
Knockdown of piRNA pathway components is associated with upregulation of *nxf2*. If *TART*-derived piRNAs are targeting *nxf2* for suppression, disruption of the piRNA pathway should relieve this suppression. We examined previously published RNA-seq data from 16 piRNA component knockdowns, as well as a control (*Yb*)(Czech et al., 2013). *Nxf2* expression increased in 14 out of 16 knockdowns, significantly more than expected by chance (one-sided binomial test P=0.002).

### Natural variation in *TART-A* copy number is correlated with *nxf2* expression levels

Previous work has shown that there is large variation in HTT element copy number at the telomeres of wild Drosophila strains (Walter et al., 2007; Wei et al., 2017). Our results predict that, if *TART-A* piRNAs are targeting *nxf2* for suppression, then strains with more copies of *TART-A* should have lower expression of *nxf2*. To test this prediction, we used previously published Illumina genomic sequencing data and microarray gene expression profiles from the Drosophila Genetic Reference Panel (DGRP)(Huang et al., 2014; Mackay et al., 2012). We used the Illumina data to infer *TART-A* copy number for 151 DGRP strains (see Methods) and obtained *nxf2* microarray gene expression levels from whole adult females for these same strains. We found that, as predicted, there is a strong negative correlation between *TART-A* copy number and *nxf2* gene expression levels among the DGRP (**Figure 7**) (Spearman’s rho = -0.48, P=4.6e-10).

**Figure 7.**
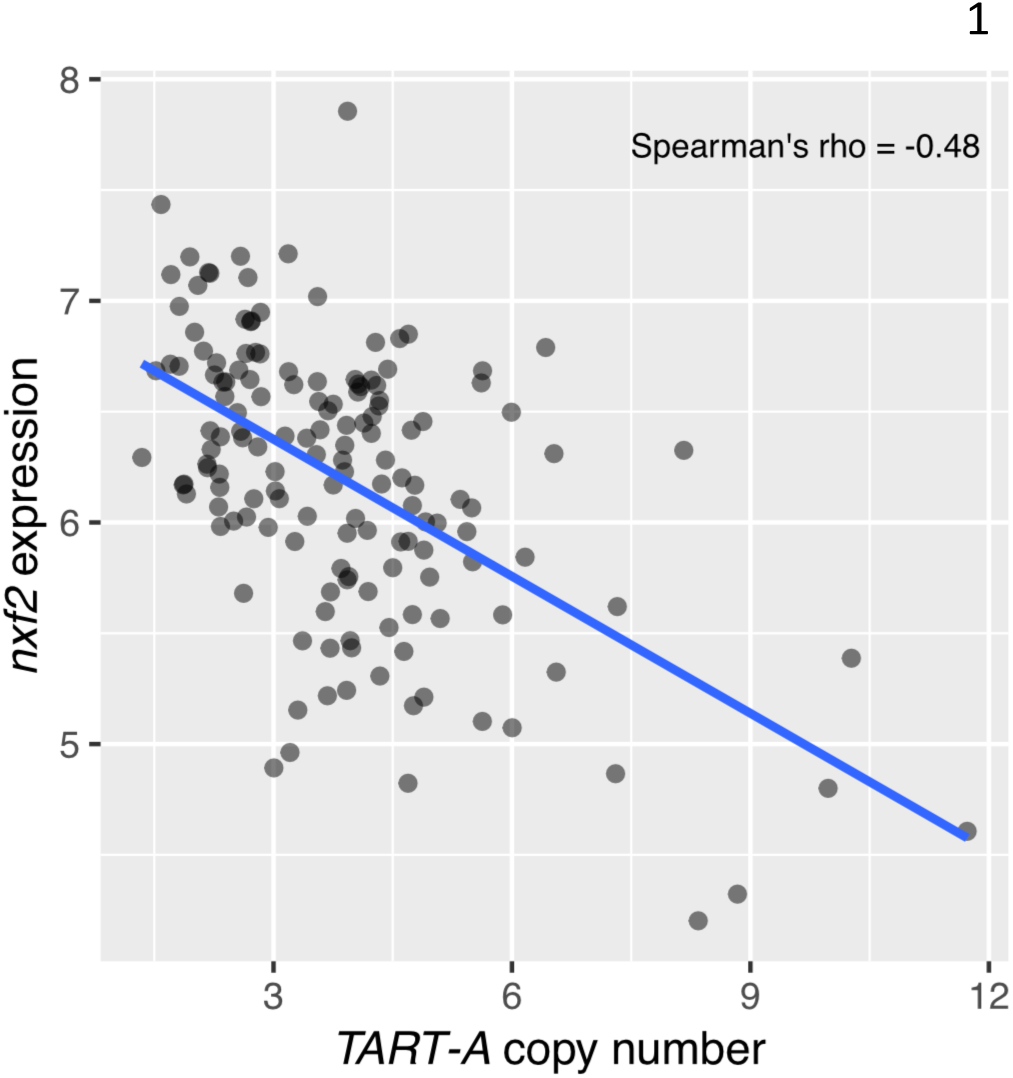
*TART-A* copy number is negatively correlated with *nxf2* expression across the Drosophila Genetic Reference Panel (DGRP). We inferred *TART-A* copy-number for 151 DGRP strains using published Illumina sequencing data (Huang et al., 2014; Mackay et al., 2012) and retrieved expression values for *nxf2* from microarray data from whole adult females (Huang et al., 2015). We found that *TART-A* copy number is significantly negatively correlated with *nxf2* expression levels, as expected if *TART-A* piRNAs are targeting *nxf2* for suppression (Spearman’s rho = -0.48, P = 4.6e-10).

## Discussion

If the coding sequence of a gene shares sequence homology with a known transposable element, the most likely explanation for this shared homology is that a portion of the gene was derived from a TE insertion. This is, understandably, what was previously reported by Sackton *et al* for the *nxf2* gene and the *TART-A* TE (Sackton et al., 2009), however our analyses are not consistent with such a scenario. Specifically, based on sequence similarity and phylogenetic clustering, the event that created the shared homology between *nxf2* and *TART-A* must have occurred relatively recently, after *D. melanogaster* diverged from *D. simulans*, yet the putative insertion of *TART-A* in the *nxf2* gene is shared across *Drosophila*. A scenario that is more consistent with these observations is one where, rather than the *nxf2* gene gaining sequence from *TART-A*, the *TART-A* element captured a portion of the *nxf2* gene, likely via aberrant transcription that extended past the internal *TART-A* poly-A signal to another poly-A signal in the flanking genomic region. This process has been observed for other TEs and is known as exon shuffling or transduction (Moran et al., 1999; Pickeral et al., 2000). Notably, the *nxf2*-like sequence of *TART-A* is located in its 3’ UTR, which would be expected if it were acquired via transduction (**Figure 1**). Interestingly, *TART* is part of the LINE family of non-LTR retrotransposons and Human LINE-L1 elements are known to undergo transduction fairly frequently (Goodier, Ostertag, & Kazazian, 2000; Moran et al., 1999; Pickeral et al., 2000). However, transduction would require that an active *TART-A* element was inserted somewhere upstream of the 3’ region of *nxf2* at some point in the *D. melanogaster* lineage, but has since been lost from the population. Is this possible given that *TART-A* should only replicate to chromosome ends? The TIDAL-fly database of polymorphic TEs in *D. melanogaster* reports several polymorphic *TART-A* insertions far from the chromosome ends, which suggests that this element is occasionally capable of inserting into locations outside of the telomeres (Rahman et al., 2015).

The aberrant *TART-A* copy that acquired a portion of the *nxf2* gene most likely arose as a single polymorphic insertion in an ancestral *D. melanogaster* population, yet the *nxf2*-like region of *TART-A* is now present in all full-length *TART-A* elements in the *D. melanogaster* reference genome assembly. We were unable to find any *D. melanogaster TART-A* elements in the reference genome, or in GenBank, whose 3’ UTR lacks the *nxf2*-like sequence. This suggests that the initially aberrant *TART-A* copy, which acquired a portion of *nxf2*, has now replaced the ancestral *TART-A* element, consistent with the gene acquisition event conferring a fitness benefit to *TART-A*.

How could the gene acquisition benefit *TART-A*? We found that the *nxf2*-like region of *TART-A* produces abundant antisense piRNAs that share homology with the *nxf2* gene, and the *nxf2* gene produces additional phased piRNAs from the unique sequence directly downstream from the regions of shared homology (**Figure 4**). These two observations are consistent with a scenario where TART-derived piRNAs guide Aub proteins to the *nxf2* transcript. The *TART-A* piRNAs may then act as “trigger” piRNAs that catalyze cleavage of *nxf2* transcripts while also resulting in the production of phased piRNAs starting in the region of shared homology and proceeding in the 3’ direction to the end of the *nxf2* transcript (**Figure 5**). The piRNA-mediated cleavage of *nxf2* transcripts, which is supported by degradome-seq data (see **Figure S5**), should result in a reduction in *nxf2* expression levels. PiRNA-mediated suppression of *nxf2* is consistent with our finding that disruption of the piRNA pathway by RNAi tends to result in increased *nxf2* expression (**Figure 6**). Given that *nxf2* plays a role in suppressing *TART-A* activity, reduced *nxf2* levels should relieve *TART-A* suppression, which would presumably increase *TART-A* fitness by allowing it to make more copies of itself. Indeed, in the DGRP, we find that individuals with lower *nxf2* expression levels tend to have higher numbers of *TART-A* copies and *vice versa* (**Figure 7**).

If additional copies of *TART-A* act to further suppress *nxf2* expression, which then further de-represses *TART-A*, why is there not run-away accumulation of telomere length in *D. melanogaster*? Previous work has shown that long telomeres in *D. melanogaster* are associated with both reduced fertility and fecundity (Walter et al., 2007), so it is possible that a run-away trend towards increasing telomere length is balanced by a fitness cost.

Targeting of host transcripts by transposon-derived piRNAs has been previously observed in Drosophila. Most notably, piRNAs from the LTR retrotransposons *roo* and *412* play a critical role in embryonic development by targeting complementary sequence in the 3’ UTR of the gene *nos*, leading to its repression in the soma (Rouget et al., 2010). More recent results suggest hundreds of maternal transcripts could be regulated in a similar fashion (Barckmann et al., 2015). However, these represent cases where TE piRNAs have been co-opted to regulate host transcripts, whereas our results suggest that the piRNA targeting of *nxf2* is a counter-defense strategy by *TART-A*. This type of strategy has only been previously observed in plants (Cosby et al., 2019). In rice, a CACTA DNA transposon produces a micro-RNA that targets a host methyltransferase gene known to be involved in TE suppression (Nosaka et al., 2012), while in Arabidopsis, siRNAs from *Athila6* retrotransposons target the stress granule protein UBP1b, which is involved in suppressing *Athila6* GAG protein production (McCue et al., 2013).

Given that viruses and other pathogens have evolved a variety of methods to block or disrupt host defense mechanisms, it is surprising that there is much less evidence for TEs adopting similar strategies (Cosby et al., 2019). However, unlike viruses, TEs depend heavily on vertical transmission from parent to offspring. Any counter-defense strategy that impacts host fitness would therefore decrease the fitness of the TE as well. Furthermore, disruption of host silencing is likely to lead to upregulation of other TEs, making it more likely that will be a severe decrease in host fitness, similar to what is observed in hybrid dysgenesis. These explanations are relevant to our results: *TART-A* may be targeting *nxf2* for its own advantage, but our knockdown experiment shows that *nxf2* suppression causes upregulation of many other TEs besides *TART-A* (**Figures 3 and S3**) and other studies have shown that *nxf2* mutants are sterile (Batki et al., 2019; Fabry et al., 2019). Why then, does *TART-A* appear to be targeting *nxf2* in spite of these potentially deleterious consequences? One possibility is that the suppression of *nxf2* expression caused by *TART-A* is relatively mild (i.e. much less than the level of down-regulation caused by the RNAi knockdown), which is enough to provide a slight benefit to *TART-A* without causing widespread TE activation. It is also possible that the suppression effect was initially much larger, but has since been counterbalanced by cis-acting variants that increase *nxf2* expression. Future work examining TE activation under varying levels of *nxf2* expression may help to determine whether there is a tipping point where *nxf2* suppression becomes catastrophic.

In summary, our results show that so-called domesticated TEs, if active, can still be in conflict with their host and raise the possibility that TE counter-defense strategies may be more common than previously recognized, despite the potentially deleterious consequences for the host.

## Methods

### *TART-A* sequence analysis

We used the *TART-A* sequence from RepBase (Jurka, 2000), which is derived from the sequence reported in (Abad et al., 2004a) (Genbank accession AJ566116). This sequence represents a single full-length *TART-A* element cloned from the *D. melanogaster* iso1 reference strain. The *nxf2*-like portion of this sequence is 100% identical to another *TART-A* element cloned and sequenced from *D. melanogaster* strain A4-4 (Genbank DMU02279)(Levis et al., 1993) as well as the *TART-A* sequence from the FlyBase canonical set of transposon sequences (version 9.42)(Thurmond et al., 2019) (cloned from *D. melanogaster* strain Oregon-R: Genbank AY561850)(Berloco, Fanti, Sheen, Levis, & Pimpinelli, 2005).

We used BLAST (Altschul et al., 1990) to compare the *TART-A* sequence to the *D. melanogaster nxf2* transcript and visualized BLAST alignments with Kablammo (Wintersinger & Wasmuth, 2015). To compare *TART-A* among Drosophila species, we used the *D. yakuba TART-A* sequence reported in (Casacuberta & Pardue, 2002)(GenBank AF468026), which includes the 3’ UTR. We also used the *D. sechellia TART-A* ORF2 reported by (Villasante et al., 2007)(Genbank AM040251) to search the *D. sechellia* FlyBase r1.3 genome assembly for a *TART-A* copy that included the 3’ UTR, which we found on scaffold_330:4944-14419. We attempted a similar approach for *D. simulans*, but were unable to find a *TART-A* copy in the *D. simulans* FlyBase r2.02 assembly that included the 3’ UTR. We aligned the *D. melanogaster, D. yakuba* and *D. sechellia TART-A* sequences to each other, and to the *D. melanogaster nxf2* transcript (FlyBase FBtr0089479), using nucmer (Kurtz et al., 2004). We then used mummerplot (Kurtz et al., 2004) to create a dotplot to visualize the alignments. To identify all copies of *TART-A* carrying the *nxf2*-like sequence, we used BLAST to search the *TART-A* 3’ UTR against the *D. melanogaster* release 6 reference genome.

### nxf2 sequence analysis

We downloaded *nxf2* transcripts from the NCBI RefSeq database for *Drosophila simulans* (XM_016169386.1), *yakuba* (XM_002095083.2), *erecta* (XM_001973010.3), *biarmipes* (XM_017111057.1), and *elegans* (XM_017273027.1) and created a codon-aware multiple sequence alignment using PRANK (Loytynoja, 2014), which we visualized with JalView (Waterhouse, Procter, Martin, Clamp, & Barton, 2009). To compare Nxf2 peptide sequences, we used the web version of NCBI BLAST to search the *D. melanogaster* Nxf2 peptide sequence against all Drosophila peptide sequences present in the RefSeq database. We then used the NCBI COBALT (Papadopoulos & Agarwala, 2007) multiple-sequence alignment tool to align the sequences shown in **Figure S1.**

### *TART-A*/*nxf2* gene tree

We extracted the *nxf2*-like sequences from all *TART-A* copies present in the *D. melanogaster* reference genome and aligned them to the *TART*-like *nxf2* sequences from seven Drosophila species using PRANK. We then inferred a maximum likelihood phylogeny with 100 bootstrap replicates using RAxML (Stamatakis, 2014).

### *nxf2* knockdown

We used two different strains from the Drosophila Transgenic RNAi Project (TRiP) that express dsRNA for RNAi of *nxf2* (Bloomington #34957 & #33985), as well as a control strain for RNAi of the *white* gene (Bloomington #33613). Seven males of each of these strains were crossed to seven, 3-5 day old, virgin females carrying the nos-GAL4 driver (Bloomington #25751). After 6 days of mating, we discarded the parental flies and then transferred F1 offspring to fresh food for 2.5 days before collecting ovaries from six females for each cross. We performed two biological replicates for each of the three crosses, dissected the ovaries in 1x PBS and immediately transferred them to RNAlater. We extracted RNA using Trizol/Phenol-Chloroform and used the AATI Fragment Analyzer to assess RNA integrity. We then prepared stranded, total RNA-seq libraries by first depleting rRNA with ribo-zero and then using the NEBnext ULTRA II library prep kit to prepare the sequencing libraries. The libraries were sequenced on the Illumina NextSeq machine with 150 bp paired-end reads.

### *nxf2* knockdown RNA-seq analysis

The average insert sizes of the total RNA-seq libraries were less than 300 bp, which resulted in overlapping mate pairs for the majority of sequenced fragments. Instead of analyzing these data as paired-end reads, we instead merged the overlapping mates to generate single-end reads using BBmerge (Bushnell, Rood, & Singer, 2017). We removed rRNA and tRNA contamination from the merged reads by aligning them to all annotated rRNA and tRNA sequences in the *D. melanogaster* reference genome using Hisat2 (Kim, Langmead, & Salzberg, 2015) and retained all unaligned reads. In order to quantify expression from genes as well as TEs, we combined all *D. melanogaster* transcript sequences (FlyBase version 6.26) with *D. melanogaster* RepBase TE consensus sequences. We accounted for multi-mapping reads by using bowtie2 (Langmead & Salzberg, 2012) to align each read to all possible alignment locations (using *--all* and *--very-sensitive-local*) and then using eXpress (Roberts & Pachter, 2013) to estimate FPKM values, accounting for the multi-mapped alignments. We averaged FPKM values between biological replicates and assessed the reproducibility of both TE and gene expression profiles in the *nxf2* knockdown by comparing the results from the two different dsRNA hairpins.

### piRNA analysis

We analyzed previously published piRNA data from 16 strains from the Drosophila Genetic Reference Panel (DGRP)(Song et al., 2014). We used cutadapt (Martin, 2011) to trim adapter sequences from each library and then removed rRNA and tRNA sequences by using bowtie (Langmead, 2010) to align the reads to all annotated rDNA and tRNA genes in the *D. melanogaster* reference genome, retaining the reads that did not align. We then created a reference database composed of the following sequence sets: a hard-masked version of the *D. melanogaster* reference genome assembly (release 6) where all TE sequences and the *nxf2* gene were replaced by N’s using RepeatMasker, the full set of *D. melanogaster* RepBase TE consensus sequences, and the *nxf2* transcript, with its *TART*-like region replaced by N’s. We used the unique-weighting mode in ShortStack (Axtell, 2013; Johnson, Yeoh, Coruh, & Axtell, 2016) to align the piRNA reads to this reference database. With this mode, ShortStack probabilistically aligns multi-mapping reads based on the abundance of uniquely mapping reads in the flanking region. We then used the ShortStack alignments and Bedtools (Quinlan & Hall, 2010) to calculate coverage for sense and antisense alignments to *TART-A* as well as *nxf2*. To test for evidence of piRNA phasing, we used the formula described in (Han et al., 2015)

### piRNA component knockdowns

We used the RNA-seq counts for *nxf2* reported in GEO accession GSE117217 from 16 RNAi knockdowns of piRNA pathway components as well as a control knockdown of the *Yb* gene (Czech et al., 2013). For each knockdown, we normalized *nxf2* expression by dividing the raw counts by the sum of all gene counts and reported the result in Reads Per Million (RPM).

### Degradome-seq analysis

We used degradome-seq and Aub-immunoprecipitated small RNA data from wild-type *D. melanogaster* strain w1 (W. Wang et al., 2014). We used bowtie2 to align the degradome-seq data to the same reference sequence used in the piRNA analysis except we unmasked the *nxf2* transcript. We analyzed the small RNA data as described under “piRNA analysis” and then used bedtools to extract degradome read alignments whose 5’ end was located in the *TART*-like region of *nxf2* and antisense small RNA alignments whose 5’ end was located in the *nxf2*-like region of *TART-A* and whose length was consistent with piRNAs (23-30 bp). We then used bowtie to align the minus strand piRNAs to the *nxf2* transcript and used bedtools to identify piRNAs whose 5’ end overlapped the 5’ of degradome reads by 10 basepairs.

### *TART-A* copy number variation and *nxf2* expression

We used Illumina genomic sequencing data from the DGRP (Huang et al., 2014; Mackay et al., 2012) to estimate *TART-A* copy number. Across strains, the DGRP Illumina data differs in terms of coverage, read length, and paired versus single-end data. To attempt to control for these differences, we trimmed all reads to 75 bp and treated all data as single-end. We also downsampled all libraries to ∼13 million reads. We first trimmed each strain’s complete dataset (unix command: *zcat file.fastq.gz* | *cut -c 75*) and then aligned the trimmed reads to the *D. melanogaster* release 6 genome assembly using bowtie2 with the *--very-sensitive* option. We then corrected the resulting bam file for GC bias using DeepTools (Ramirez, Dundar, Diehl, Gruning, & Manke, 2014) and counted the number of aligned reads in the corrected bam file using samtools (Li et al., 2009). We removed all strains with less than 13 million aligned reads and, for each remaining strain, we calculated the fraction of reads to keep by dividing the smallest number of aligned reads across all remaining individuals (13,594,737) by the total number of aligned reads for that strain. We then used this fraction to randomly downsample the GC corrected bam file using the *subsample* option from *samtools view* (Li et al., 2009). We converted each bam file to a fastq file with *samtools fastq* and aligned the fastq file to the *D. melanogaster* RepBase TE sequences with *bowtie2* using the *--very-sensitive, --local*, and *--all* options. With --*all, bowtie2* reports every possible alignment for each multi-mapping sequence. We then used *eXpress* to retain a single alignment for each multi-mapping sequence based on the abundance of neighboring unique alignments. We used the *eXpress* bam files to calculate the median per-base coverage (excluding positions with coverage of zero) for the *TART-A* coding sequence (i.e. ORF1 & ORF2), for each individual. To estimate *TART-A* copy number, we divided the median *TART-A* coverage of each strain by that strain’s median per-base coverage of all uniquely-mappable positions in the *D. melanogaster* reference genome (calculated from the GC corrected, downsampled bam file). Uniquely-mappable positions were identified using *mirth* (https://github.com/EvolBioInf/mirth). We obtained *nxf2* expression values from previously published microarray gene expression profiles from whole adult females for all DGRP strains (Huang et al., 2015).

## Supplemental Figures

**Figure S1.**
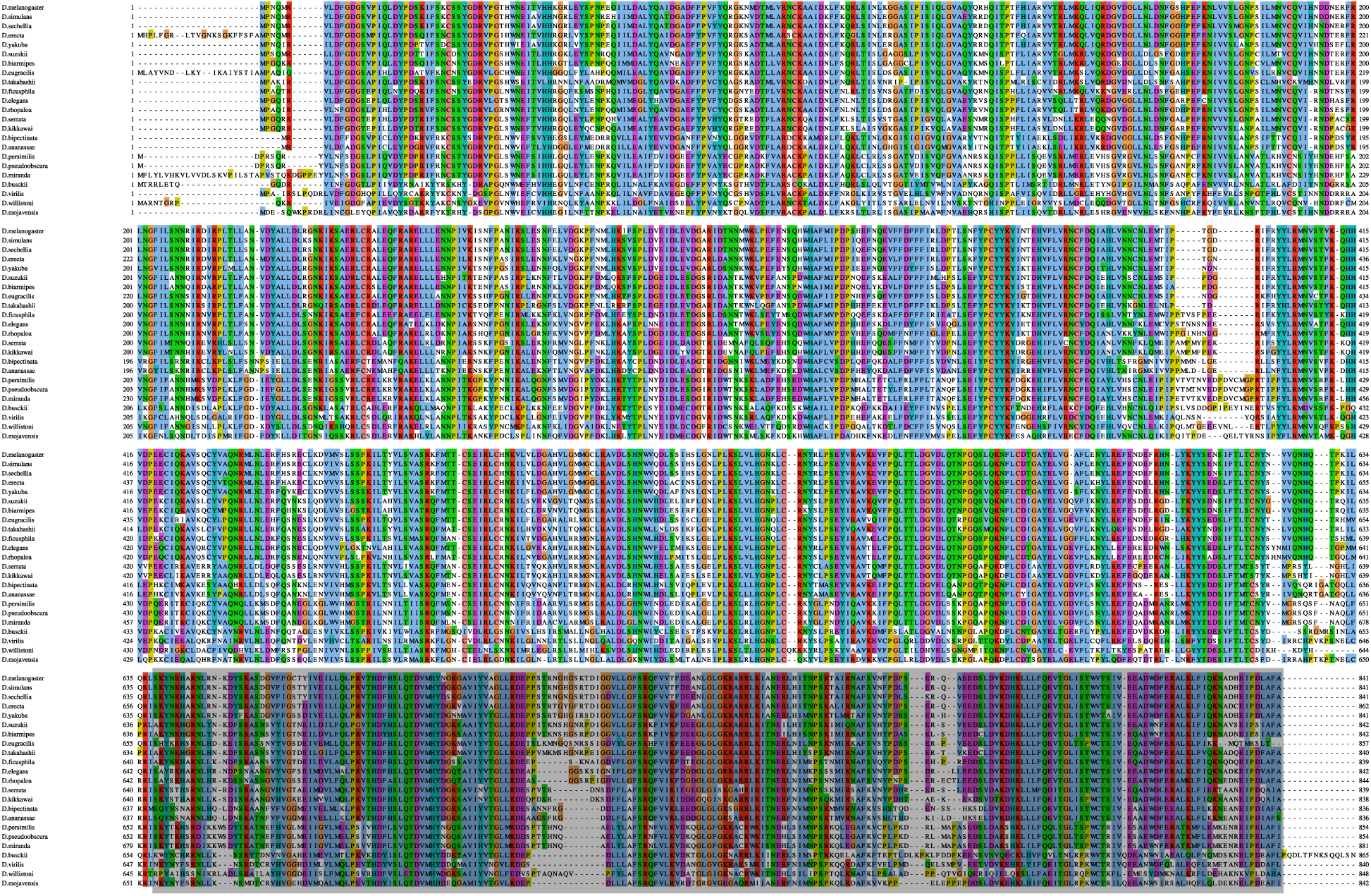
Peptide alignment of Nxf2 homologs. We used NCBI web BLAST to search the *D. melanogaster* Nxf2 peptide sequence against the RefSeq peptide database and identified homologs in 22 Drosophila species. The C-terminal region of Nxf2 derives from coding sequence which shares homology with the *TART-A* transposable element (grey box). At the peptide level, this region is conserved out to *D. virilis*, which suggests that, if it was acquired from an insertion of the *TART-A* TE, the insertion would have occurred in the common ancestor of the entire genus.

**Figure S2.**
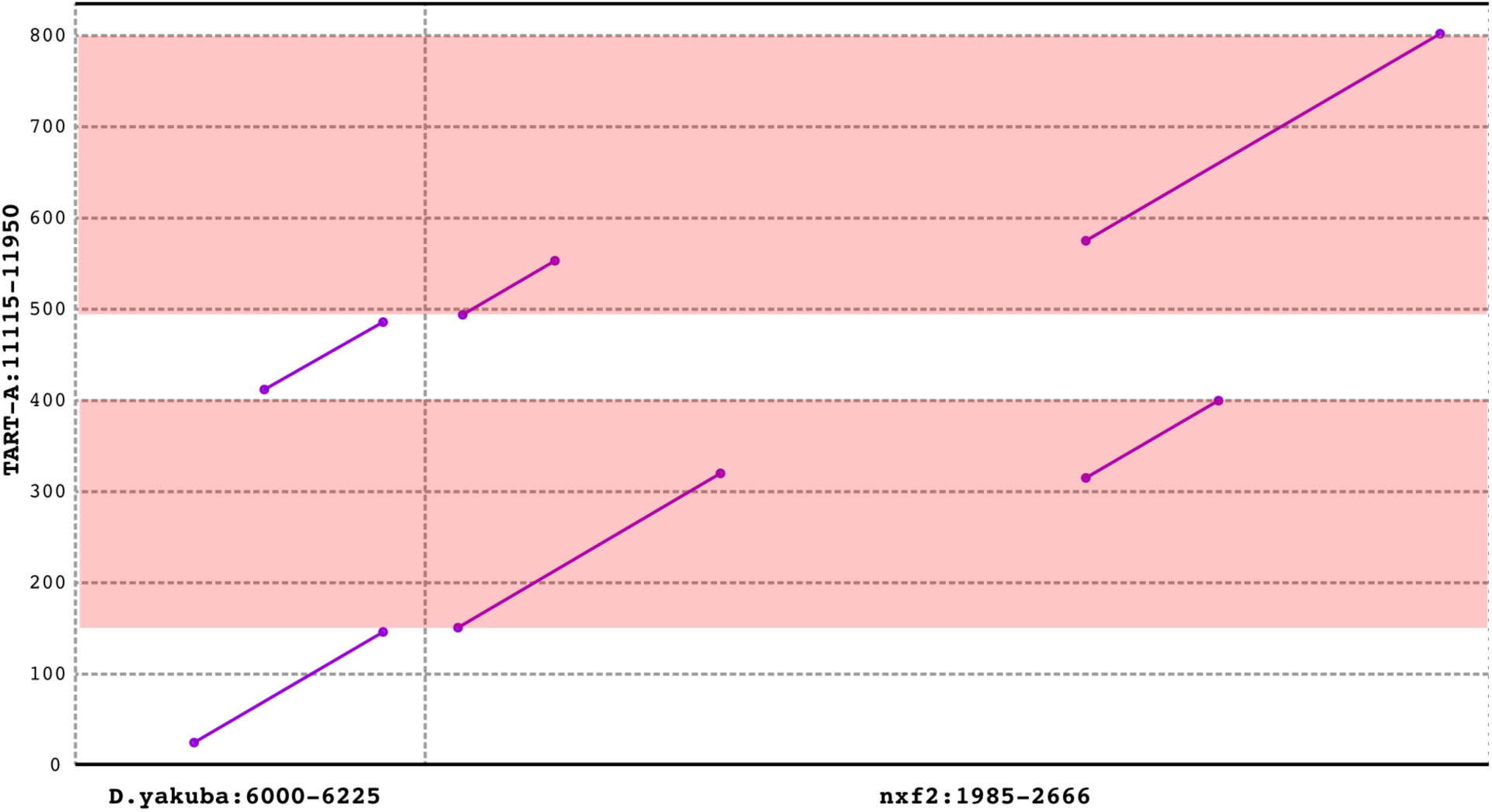
Zoom view of dotplot showing alignments of *D. melanogaster TART-A* versus *D. melanogaster nxf2* and *D. yakuba TART-A*. The pink boxes show the two segments of shared homology between *D. melanogaster TART-A* and *D. melanogaster nxf2. D. yakuba TART-A* aligns to *D. melanogaster TART-A* at regions directly adjacent to, but not including, the *TART-A/nxf2* shared homology.

**Figure S3.**
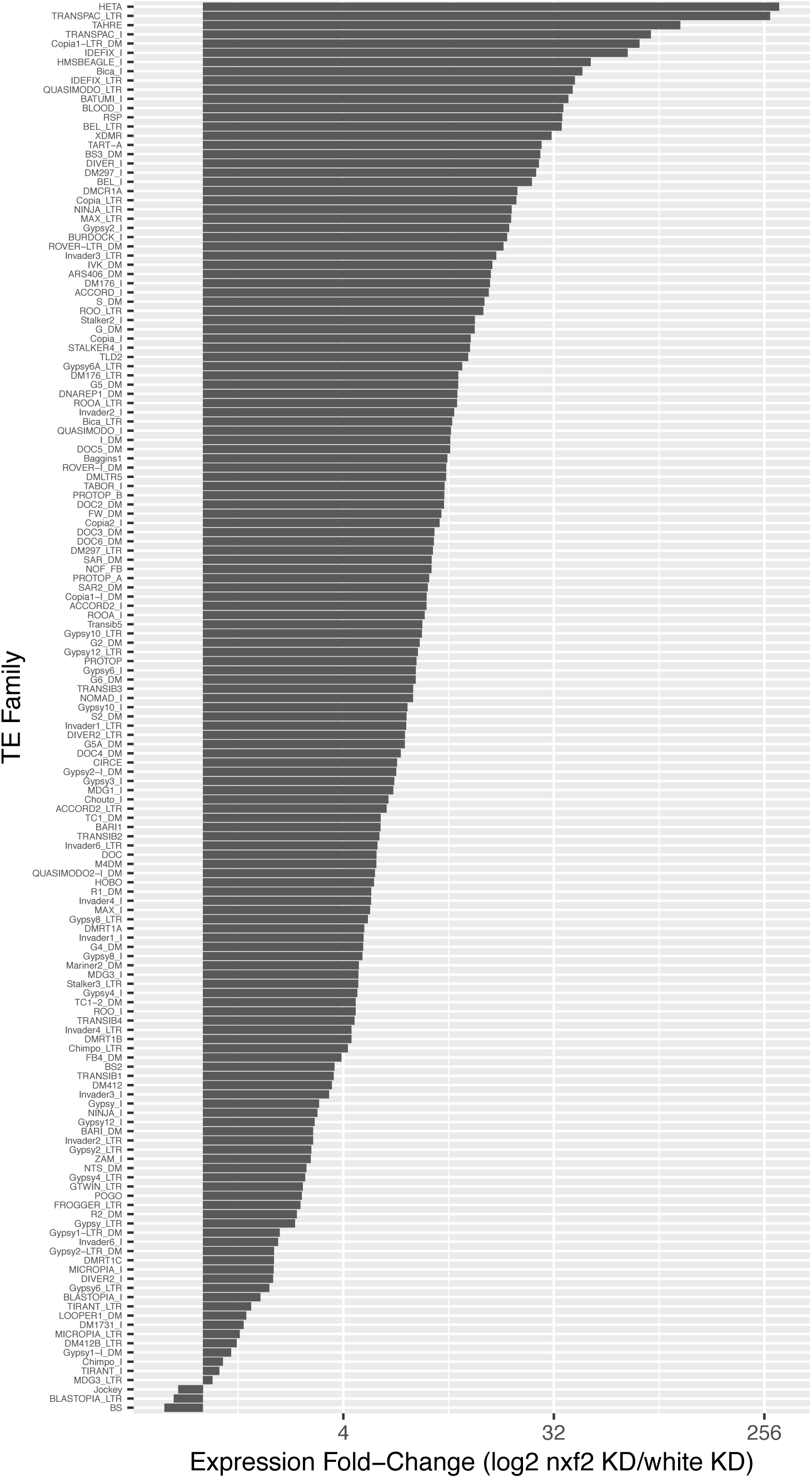
Repetitive element upregulation in *nxf2* knockdown. Each RepBase repeat for which we observed expression in total RNA-seq data from female ovaries is shown on the y-axis and the fold-change in expression in the *nxf2* RNAi knockdown versus a control knockdown of the *white* gene is shown on the x-axis with a log2 scale. Expression values are the mean of two biological replicates for both knockdown and control. For LTR retrotransposons, LTRs are shown separately from the rest of the TE.

**Figure S4.**
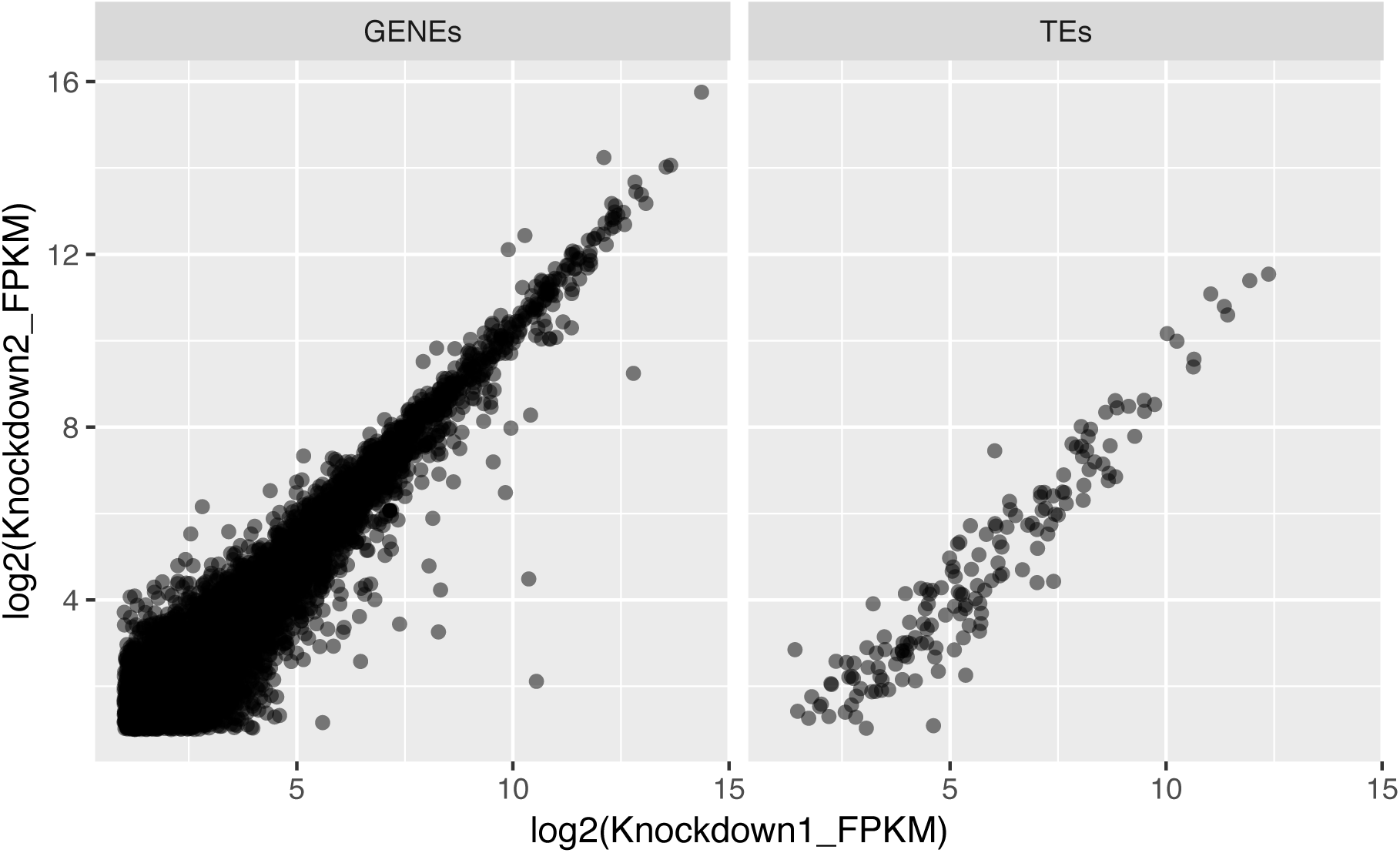
Correlation between shRNAs in *nxf2* knockdown. We used two shRNAs that target different regions of the *nxf2* transcript and calculated expression values for genes as well as TEs for each knockdown. We found that the expression values are highly correlated between the two experiments (Spearman’s rho=0.92 [Genes] and 0.94 [TEs]).

**Figure S5.**
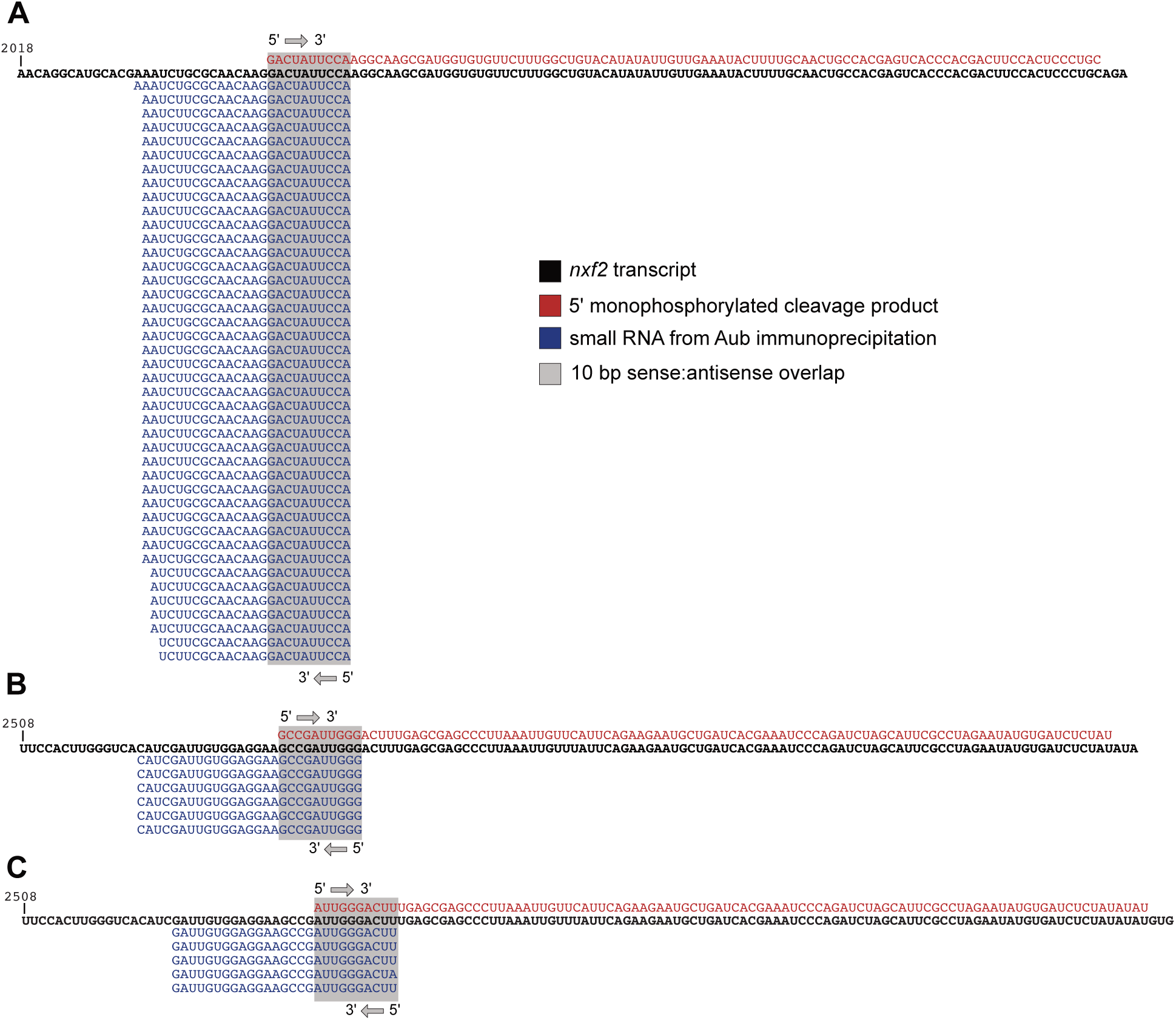
*nxf2* cleavage products from degradome-seq data. We analyzed published degradome-seq and Aub-immunoprecipitated small RNA data to determine whether there were *nxf2* degradome-seq reads showing the 10 bp sense:antisense overlap with *TART-A* piRNAs, consistent with cleavage by a piwi protein. We identified three locations (A – C above) within the *TART*-like region of *nxf2* where degradome-seq cleavage products (red) overlap with antisense piRNAs (blue) by 10 bp at their 5’ ends. The *nxf2* transcript is shown in black.

## References

Abad, J. P., De Pablos, B., Osoegawa, K., De Jong, P. J., Martin-Gallardo, A., & Villasante, A. (2004a). Genomic analysis of Drosophila melanogaster telomeres: full-length copies of HeT-A and TART elements at telomeres. Mol Biol Evol, 21 (9), 1613–1619. doi:10.1093/molbev/msh174

Abad, J. P., De Pablos, B., Osoegawa, K., De Jong, P. J., Martin-Gallardo, A., & Villasante, A. (2004b). TAHRE, a novel telomeric retrotransposon from Drosophila melanogaster, reveals the origin of Drosophila telomeres. Mol Biol Evol, 21 (9), 1620–1624. doi:10.1093/molbev/msh180

Addo-Quaye, C., Eshoo, T. W., Bartel, D. P., & Axtell, M. J. (2008). Endogenous siRNA and miRNA targets identified by sequencing of the Arabidopsis degradome. Curr Biol, 18 (10), 758–762. doi:10.1016/j.cub.2008.04.042

Altschul, S. F., Gish, W., Miller, W., Myers, E. W., & Lipman, D. J. (1990). Basic local alignment search tool. J Mol Biol, 215 (3), 403–410. doi:10.1016/S0022-2836(05)80360-2

Axtell, M. J. (2013). ShortStack: comprehensive annotation and quantification of small RNA genes. RNA, 19 (6), 740–751. doi:10.1261/rna.035279.112

Barckmann, B., Pierson, S., Dufourt, J., Papin, C., Armenise, C., Port, F., … Simonelig, M. (2015). Aubergine iCLIP Reveals piRNA-Dependent Decay of mRNAs Involved in Germ Cell Development in the Early Embryo. Cell Rep, 12 (7), 1205–1216. doi:10.1016/j.celrep.2015.07.030

Batki, J., Schnabl, J., Wang, J., Handler, D., Andreev, V. I., Stieger, C. E., … Brennecke, J. (2019). The nascent RNA binding complex SFiNX licenses piRNA-guided heterochromatin formation. Nat Struct Mol Biol, 26 (8), 720–731. doi:10.1038/s41594-019-0270-6

Berloco, M., Fanti, L., Sheen, F., Levis, R. W., & Pimpinelli, S. (2005). Heterochromatic distribution of HeT-A- and TART-like sequences in several Drosophila species. Cytogenet Genome Res, 110 (1-4), 124–133. doi:10.1159/000084944

Biessmann, H., & Mason, J. M. (1997). Telomere maintenance without telomerase. Chromosoma, 106 (2), 63–69.

Biessmann, H., Valgeirsdottir, K., Lofsky, A., Chin, C., Ginther, B., Levis, R. W., & Pardue, M. L. (1992). HeT-A, a transposable element specifically involved in “healing” broken chromosome ends in Drosophila melanogaster. Mol Cell Biol, 12 (9), 3910–3918. doi:10.1128/mcb.12.9.3910

Blumenstiel, J. P., Erwin, A. A., & Hemmer, L. W. (2016). What Drives Positive Selection in the Drosophila piRNA Machinery? The Genomic Autoimmunity Hypothesis. Yale J Biol Med, 89 (4), 499–512.

Bohne, A., Brunet, F., Galiana-Arnoux, D., Schultheis, C., & Volff, J. N. (2008). Transposable elements as drivers of genomic and biological diversity in vertebrates. Chromosome Res, 16 (1), 203–215. doi:10.1007/s10577-007-1202-6

Brennecke, J., Aravin, A. A., Stark, A., Dus, M., Kellis, M., Sachidanandam, R., & Hannon, G. J. (2007). Discrete small RNA-generating loci as master regulators of transposon activity in Drosophila. Cell, 128 (6), 1089–1103. doi:10.1016/j.cell.2007.01.043

Bushnell, B., Rood, J., & Singer, E. (2017). BBMerge - Accurate paired shotgun read merging via overlap. PLoS One, 12 (10), e0185056. doi:10.1371/journal.pone.0185056

Cam, H. P., Noma, K., Ebina, H., Levin, H. L., & Grewal, S. I. (2008). Host genome surveillance for retrotransposons by transposon-derived proteins. Nature, 451 (7177), 431–436. doi:10.1038/nature06499

Casacuberta, E., & Pardue, M. L. (2002). Coevolution of the telomeric retrotransposons across Drosophila species. Genetics, 161 (3), 1113–1124.

Chang, C. H., Chavan, A., Palladino, J., Wei, X., Martins, N. M. C., Santinello, B., … Mellone, B. G. (2019). Islands of retroelements are major components of Drosophila centromeres. PLoS Biol, 17 (5), e3000241. doi:10.1371/journal.pbio.3000241

Chueh, A. C., Northrop, E. L., Brettingham-Moore, K. H., Choo, K. H., & Wong, L. H. (2009). LINE retrotransposon RNA is an essential structural and functional epigenetic component of a core neocentromeric chromatin. PLoS Genet, 5 (1), e1000354. doi:10.1371/journal.pgen.1000354

Chuong, E. B., Elde, N. C., & Feschotte, C. (2016). Regulatory evolution of innate immunity through cooption of endogenous retroviruses. Science, 351 (6277), 1083–1087. doi:10.1126/science.aad5497

Chuong, E. B., Elde, N. C., & Feschotte, C. (2017). Regulatory activities of transposable elements: from conflicts to benefits. Nat Rev Genet, 18 (2), 71–86. doi:10.1038/nrg.2016.139

Chuong, E. B., Rumi, M. A., Soares, M. J., & Baker, J. C. (2013). Endogenous retroviruses function as species-specific enhancer elements in the placenta. Nat Genet, 45 (3), 325–329. doi:10.1038/ng.2553

Cosby, R. L., Chang, N. C., & Feschotte, C. (2019). Host-transposon interactions: conflict, cooperation, and cooption. Genes Dev, 33 (17-18), 1098–1116. doi:10.1101/gad.327312.119

Crysnanto, D., & Obbard, D. J. (2019). Widespread gene duplication and adaptive evolution in the RNA interference pathways of the Drosophila obscura group. BMC Evol Biol, 19 (1), 99. doi:10.1186/s12862-019-1425-0

Czech, B., Preall, J. B., McGinn, J., & Hannon, G. J. (2013). A transcriptome-wide RNAi screen in the Drosophila ovary reveals factors of the germline piRNA pathway. Mol Cell, 50 (5), 749–761. doi:10.1016/j.molcel.2013.04.007

Danilevskaya, O. N., Tan, C., Wong, J., Alibhai, M., & Pardue, M. L. (1998). Unusual features of the Drosophila melanogaster telomere transposable element HeT-A are conserved in Drosophila yakuba telomere elements. Proc Natl Acad Sci U S A, 95 (7), 3770–3775. doi:10.1073/pnas.95.7.3770

Dennis, C., Brasset, E., Sarkar, A., & Vaury, C. (2016). Export of piRNA precursors by EJC triggers assembly of cytoplasmic Yb-body in Drosophila. Nat Commun, 7, 13739. doi:10.1038/ncomms13739

Dunn-Fletcher, C. E., Muglia, L. M., Pavlicev, M., Wolf, G., Sun, M. A., Hu, Y. C., … Muglia, L. J. (2018). Anthropoid primate-specific retroviral element THE1B controls expression of CRH in placenta and alters gestation length. PLoS Biol, 16 (9), e2006337. doi:10.1371/journal.pbio.2006337

Ellison, C., & Bachtrog, D. (2019). Contingency in the convergent evolution of a regulatory network: Dosage compensation in Drosophila. PLoS Biol, 17 (2), e3000094. doi:10.1371/journal.pbio.3000094

Ellison, C. E., & Bachtrog, D. (2013). Dosage compensation via transposable element mediated rewiring of a regulatory network. Science, 342 (6160), 846–850. doi:10.1126/science.1239552

Esnault, C., Heidmann, O., Delebecque, F., Dewannieux, M., Ribet, D., Hance, A. J., … Schwartz, O. (2005). APOBEC3G cytidine deaminase inhibits retrotransposition of endogenous retroviruses. Nature, 433 (7024), 430–433. doi:10.1038/nature03238

Fabry, M. H., Ciabrelli, F., Munafo, M., Eastwood, E. L., Kneuss, E., Falciatori, I., … Czech, B. (2019). piRNA-guided co-transcriptional silencing coopts nuclear export factors. Elife, 8. doi:10.7554/eLife.47999

Feschotte, C. (2008). Transposable elements and the evolution of regulatory networks. Nat Rev Genet, 9 (5), 397–405. doi:10.1038/nrg2337

Fu, Y., Kawabe, A., Etcheverry, M., Ito, T., Toyoda, A., Fujiyama, A., … Kakutani, T. (2013). Mobilization of a plant transposon by expression of the transposon-encoded anti-silencing factor. EMBO J, 32 (17), 2407–2417. doi:10.1038/emboj.2013.169

Fuentes, D. R., Swigut, T., & Wysocka, J. (2018). Systematic perturbation of retroviral LTRs reveals widespread long-range effects on human gene regulation. Elife, 7. doi:10.7554/eLife.35989

George, J. A., Traverse, K. L., DeBaryshe, P. G., Kelley, K. J., & Pardue, M. L. (2010). Evolution of diverse mechanisms for protecting chromosome ends by Drosophila TART telomere retrotransposons. Proc Natl Acad Sci U S A, 107 (49), 21052–21057. doi:10.1073/pnas.1015926107

Goodier, J. L., Ostertag, E. M., & Kazazian, H. H., Jr. (2000). Transduction of 3’-flanking sequences is common in L1 retrotransposition. Hum Mol Genet, 9 (4), 653–657. doi:10.1093/hmg/9.4.653

Gunawardane, L. S., Saito, K., Nishida, K. M., Miyoshi, K., Kawamura, Y., Nagami, T., … Siomi, M. C. (2007). A slicer-mediated mechanism for repeat-associated siRNA 5’ end formation in Drosophila. Science, 315 (5818), 1587–1590. doi:10.1126/science.1140494

Han, B. W., Wang, W., Li, C., Weng, Z., & Zamore, P. D. (2015). Noncoding RNA. piRNA-guided transposon cleavage initiates Zucchini-dependent, phased piRNA production. Science, 348 (6236), 817–821. doi:10.1126/science.aaa1264

Helleu, Q., & Levine, M. T. (2018). Recurrent Amplification of the Heterochromatin Protein 1 (HP1) Gene Family across Diptera. Mol Biol Evol, 35 (10), 2375–2389. doi:10.1093/molbev/msy128

Herold, A., Suyama, M., Rodrigues, J. P., Braun, I. C., Kutay, U., Carmo-Fonseca, M., … Izaurralde, E. (2000). TAP (NXF1) belongs to a multigene family of putative RNA export factors with a conserved modular architecture. Mol Cell Biol, 20 (23), 8996–9008. doi:10.1128/mcb.20.23.8996-9008.2000

Huang, W., Carbone, M. A., Magwire, M. M., Peiffer, J. A., Lyman, R. F., Stone, E. A., … Mackay, T. F. (2015). Genetic basis of transcriptome diversity in Drosophila melanogaster. Proc Natl Acad Sci U S A, 112 (44), E6010–6019. doi:10.1073/pnas.1519159112

Huang, W., Massouras, A., Inoue, Y., Peiffer, J., Ramia, M., Tarone, A. M., … Mackay, T. F. (2014). Natural variation in genome architecture among 205 Drosophila melanogaster Genetic Reference Panel lines. Genome Res, 24 (7), 1193–1208. doi:10.1101/gr.171546.113

Hur, J. K., Luo, Y., Moon, S., Ninova, M., Marinov, G. K., Chung, Y. D., & Aravin, A. A. (2016). Splicing-independent loading of TREX on nascent RNA is required for efficient expression of dual-strand piRNA clusters in Drosophila. Genes Dev, 30 (7), 840–855. doi:10.1101/gad.276030.115

Jacobs, F. M., Greenberg, D., Nguyen, N., Haeussler, M., Ewing, A. D., Katzman, S., … Haussler, D. (2014). An evolutionary arms race between KRAB zinc-finger genes ZNF91/93 and SVA/L1 retrotransposons. Nature, 516 (7530), 242–245. doi:10.1038/nature13760

Johnson, N. R., Yeoh, J. M., Coruh, C., & Axtell, M. J. (2016). Improved Placement of Multi-mapping Small RNAs. G3 (Bethesda), 6 (7), 2103–2111. doi:10.1534/g3.116.030452

Joly-Lopez, Z., & Bureau, T. E. (2018). Exaptation of transposable element coding sequences. Curr Opin Genet Dev, 49, 34–42. doi:10.1016/j.gde.2018.02.011

Joly-Lopez, Z., Hoen, D. R., Blanchette, M., & Bureau, T. E. (2016). Phylogenetic and Genomic Analyses Resolve the Origin of Important Plant Genes Derived from Transposable Elements. Mol Biol Evol, 33 (8), 1937–1956. doi:10.1093/molbev/msw067

Jurka, J. (2000). Repbase update: a database and an electronic journal of repetitive elements. Trends Genet, 16 (9), 418–420. doi:10.1016/s0168-9525(00)02093-x

Kapusta, A., Kronenberg, Z., Lynch, V. J., Zhuo, X., Ramsay, L., Bourque, G., … Feschotte, C. (2013). Transposable elements are major contributors to the origin, diversification, and regulation of vertebrate long noncoding RNAs. PLoS Genet, 9 (4), e1003470. doi:10.1371/journal.pgen.1003470

Kelleher, E. S., Edelman, N. B., & Barbash, D. A. (2012). Drosophila interspecific hybrids phenocopy piRNA-pathway mutants. PLoS Biol, 10 (11), e1001428. doi:10.1371/journal.pbio.1001428

Khurana, J. S., Xu, J., Weng, Z., & Theurkauf, W. E. (2010). Distinct functions for the Drosophila piRNA pathway in genome maintenance and telomere protection. PLoS Genet, 6 (12), e1001246. doi:10.1371/journal.pgen.1001246

Kim, D., Langmead, B., & Salzberg, S. L. (2015). HISAT: a fast spliced aligner with low memory requirements. Nat Methods, 12 (4), 357–360. doi:10.1038/nmeth.3317

Klein, S. J., & O’Neill, R. J. (2018). Transposable elements: genome innovation, chromosome diversity, and centromere conflict. Chromosome Res, 26 (1-2), 5–23. doi:10.1007/s10577-017-9569-5

Kolaczkowski, B., Hupalo, D. N., & Kern, A. D. (2011). Recurrent adaptation in RNA interference genes across the Drosophila phylogeny. Mol Biol Evol, 28 (2), 1033–1042. doi:10.1093/molbev/msq284

Kurtz, S., Phillippy, A., Delcher, A. L., Smoot, M., Shumway, M., Antonescu, C., & Salzberg, S. L. (2004). Versatile and open software for comparing large genomes. Genome Biol, 5 (2), R12. doi:10.1186/gb-2004-5-2-r12

Langmead, B. (2010). Aligning short sequencing reads with Bowtie. Curr Protoc Bioinformatics, Chapter 11, Unit 11 17. doi:10.1002/0471250953.bi1107s32

Langmead, B., & Salzberg, S. L. (2012). Fast gapped-read alignment with Bowtie 2. Nat Methods, 9(4), 357–359. doi:10.1038/nmeth.1923

Lee, Y. C. (2015). The Role of piRNA-Mediated Epigenetic Silencing in the Population Dynamics of Transposable Elements in Drosophila melanogaster. PLoS Genet, 11 (6), e1005269. doi:10.1371/journal.pgen.1005269

Lee, Y. C., Leek, C., & Levine, M. T. (2017). Recurrent Innovation at Genes Required for Telomere Integrity in Drosophila. Mol Biol Evol, 34 (2), 467–482. doi:10.1093/molbev/msw248

Lee, Y. C. G., & Karpen, G. H. (2017). Pervasive epigenetic effects of Drosophila euchromatic transposable elements impact their evolution. Elife, 6. doi:10.7554/eLife.25762

Levine, M. T., Vander Wende, H. M., Hsieh, E., Baker, E. P., & Malik, H. S. (2016). Recurrent Gene Duplication Diversifies Genome Defense Repertoire in Drosophila. Mol Biol Evol, 33 (7), 1641–1653. doi:10.1093/molbev/msw053

Levis, R. W., Ganesan, R., Houtchens, K., Tolar, L. A., & Sheen, F. M. (1993). Transposons in place of telomeric repeats at a Drosophila telomere. Cell, 75 (6), 1083–1093. doi:10.1016/0092-8674(93)90318-k

Li, H., Handsaker, B., Wysoker, A., Fennell, T., Ruan, J., Homer, N., … Genome Project Data Processing, S. (2009). The Sequence Alignment/Map format and SAMtools. Bioinformatics, 25 (16), 2078–2079. doi:10.1093/bioinformatics/btp352

Loytynoja, A. (2014). Phylogeny-aware alignment with PRANK. Methods Mol Biol, 1079, 155–170. doi:10.1007/978-1-62703-646-7_10

Lynch, V. J., Leclerc, R. D., May, G., & Wagner, G. P. (2011). Transposon-mediated rewiring of gene regulatory networks contributed to the evolution of pregnancy in mammals. Nat Genet, 43 (11), 1154–1159. doi:10.1038/ng.917

Lynch, V. J., Nnamani, M. C., Kapusta, A., Brayer, K., Plaza, S. L., Mazur, E. C., … Wagner, G. P. (2015). Ancient transposable elements transformed the uterine regulatory landscape and transcriptome during the evolution of mammalian pregnancy. Cell Rep, 10 (4), 551–561. doi:10.1016/j.celrep.2014.12.052

Mackay, T. F., Richards, S., Stone, E. A., Barbadilla, A., Ayroles, J. F., Zhu, D., … Gibbs, R. A. (2012). The Drosophila melanogaster Genetic Reference Panel. Nature, 482 (7384), 173–178. doi:10.1038/nature10811

Malik, H. S., Burke, W. D., & Eickbush, T. H. (1999). The age and evolution of non-LTR retrotransposable elements. Mol Biol Evol, 16 (6), 793–805. doi:10.1093/oxfordjournals.molbev.a026164

Mari-Ordonez, A., Marchais, A., Etcheverry, M., Martin, A., Colot, V., & Voinnet, O. (2013). Reconstructing de novo silencing of an active plant retrotransposon. Nat Genet, 45 (9), 1029–1039. doi:10.1038/ng.2703

Martin, M. (2011). Cutadapt removes adapter sequences from high-throughput sequencing reads. 2011, 17 (1), 3. doi:10.14806/ej.17.1.200

McCue, A. D., Nuthikattu, S., & Slotkin, R. K. (2013). Genome-wide identification of genes regulated in trans by transposable element small interfering RNAs. RNA Biol, 10 (8), 1379–1395. doi:10.4161/rna.25555

Mohn, F., Handler, D., & Brennecke, J. (2015). Noncoding RNA. piRNA-guided slicing specifies transcripts for Zucchini-dependent, phased piRNA biogenesis. Science, 348 (6236), 812–817. doi:10.1126/science.aaa1039

Molaro, A., & Malik, H. S. (2016). Hide and seek: how chromatin-based pathways silence retroelements in the mammalian germline. Curr Opin Genet Dev, 37, 51–58. doi:10.1016/j.gde.2015.12.001

Moran, J. V., DeBerardinis, R. J., & Kazazian, H. H., Jr. (1999). Exon shuffling by L1 retrotransposition. Science, 283 (5407), 1530–1534. doi:10.1126/science.283.5407.1530

Murano, K., Iwasaki, Y. W., Ishizu, H., Mashiko, A., Shibuya, A., Kondo, S., … Siomi, H. (2019). Nuclear RNA export factor variant initiates piRNA-guided co-transcriptional silencing. EMBO J, 38 (17), e102870. doi:10.15252/embj.2019102870

Nosaka, M., Itoh, J., Nagato, Y., Ono, A., Ishiwata, A., & Sato, Y. (2012). Role of transposon-derived small RNAs in the interplay between genomes and parasitic DNA in rice. PLoS Genet, 8 (9), e1002953. doi:10.1371/journal.pgen.1002953

Notwell, J. H., Chung, T., Heavner, W., & Bejerano, G. (2015). A family of transposable elements co-opted into developmental enhancers in the mouse neocortex. Nat Commun, 6, 6644. doi:10.1038/ncomms7644

Obbard, D. J., Jiggins, F. M., Bradshaw, N. J., & Little, T. J. (2011). Recent and recurrent selective sweeps of the antiviral RNAi gene Argonaute-2 in three species of Drosophila. Mol Biol Evol, 28 (2), 1043–1056. doi:10.1093/molbev/msq280

Obbard, D. J., Jiggins, F. M., Halligan, D. L., & Little, T. J. (2006). Natural selection drives extremely rapid evolution in antiviral RNAi genes. Curr Biol, 16 (6), 580–585. doi:10.1016/j.cub.2006.01.065

Obbard, D. J., Maclennan, J., Kim, K. W., Rambaut, A., O’Grady, P. M., & Jiggins, F. M. (2012). Estimating divergence dates and substitution rates in the Drosophila phylogeny. Mol Biol Evol, 29 (11), 3459–3473. doi:10.1093/molbev/mss150

Papadopoulos, J. S., & Agarwala, R. (2007). COBALT: constraint-based alignment tool for multiple protein sequences. Bioinformatics, 23 (9), 1073–1079. doi:10.1093/bioinformatics/btm076

Parhad, S. S., & Theurkauf, W. E. (2019). Rapid evolution and conserved function of the piRNA pathway. Open Biol, 9 (1), 180181. doi:10.1098/rsob.180181

Petrov, D. A., Fiston-Lavier, A. S., Lipatov, M., Lenkov, K., & Gonzalez, J. (2011). Population genomics of transposable elements in Drosophila melanogaster. Mol Biol Evol, 28 (5), 1633–1644. doi:10.1093/molbev/msq337

Pickeral, O. K., Makalowski, W., Boguski, M. S., & Boeke, J. D. (2000). Frequent human genomic DNA transduction driven by LINE-1 retrotransposition. Genome Res, 10 (4), 411–415. doi:10.1101/gr.10.4.411

Pontis, J., Planet, E., Offner, S., Turelli, P., Duc, J., Coudray, A., … Trono, D. (2019). Hominoid-Specific Transposable Elements and KZFPs Facilitate Human Embryonic Genome Activation and Control Transcription in Naive Human ESCs. Cell Stem Cell, 24 (5), 724–735 e725. doi:10.1016/j.stem.2019.03.012

Quinlan, A. R., & Hall, I. M. (2010). BEDTools: a flexible suite of utilities for comparing genomic features. Bioinformatics, 26 (6), 841–842. doi:10.1093/bioinformatics/btq033

Rahman, R., Chirn, G. W., Kanodia, A., Sytnikova, Y. A., Brembs, B., Bergman, C. M., & Lau, N. C. (2015). Unique transposon landscapes are pervasive across Drosophila melanogaster genomes. Nucleic Acids Res, 43 (22), 10655–10672. doi:10.1093/nar/gkv1193

Ramirez, F., Dundar, F., Diehl, S., Gruning, B. A., & Manke, T. (2014). deepTools: a flexible platform for exploring deep-sequencing data. Nucleic Acids Res, 42(Web Server issue), W187–191. doi:10.1093/nar/gku365

Roberts, A., & Pachter, L. (2013). Streaming fragment assignment for real-time analysis of sequencing experiments. Nat Methods, 10 (1), 71–73. doi:10.1038/nmeth.2251

Rouget, C., Papin, C., Boureux, A., Meunier, A. C., Franco, B., Robine, N., … Simonelig, M. (2010). Maternal mRNA deadenylation and decay by the piRNA pathway in the early Drosophila embryo. Nature, 467 (7319), 1128–1132. doi:10.1038/nature09465

Sackton, T. B., Kulathinal, R. J., Bergman, C. M., Quinlan, A. R., Dopman, E. B., Carneiro, M., … Clark, A. G. (2009). Population genomic inferences from sparse high-throughput sequencing of two populations of Drosophila melanogaster. Genome Biol Evol, 1, 449–465. doi:10.1093/gbe/evp048

Saint-Leandre, B., Nguyen, S. C., & Levine, M. T. (2019). Diversification and collapse of a telomere elongation mechanism. Genome Res, 29 (6), 920–931. doi:10.1101/gr.245001.118

Satyaki, P. R., Cuykendall, T. N., Wei, K. H., Brideau, N. J., Kwak, H., Aruna, S., … Barbash, D. A. (2014). The Hmr and Lhr hybrid incompatibility genes suppress a broad range of heterochromatic repeats. PLoS Genet, 10 (3), e1004240. doi:10.1371/journal.pgen.1004240

Savitsky, M., Kravchuk, O., Melnikova, L., & Georgiev, P. (2002). Heterochromatin protein 1 is involved in control of telomere elongation in Drosophila melanogaster. Mol Cell Biol, 22 (9), 3204–3218. doi:10.1128/mcb.22.9.3204-3218.2002

Savitsky, M., Kwon, D., Georgiev, P., Kalmykova, A., & Gvozdev, V. (2006). Telomere elongation is under the control of the RNAi-based mechanism in the Drosophila germline. Genes Dev, 20 (3), 345–354. doi:10.1101/gad.370206

Sheen, F. M., & Levis, R. W. (1994). Transposition of the LINE-like retrotransposon TART to Drosophila chromosome termini. Proc Natl Acad Sci U S A, 91 (26), 12510–12514. doi:10.1073/pnas.91.26.12510

Shpiz, S., & Kalmykova, A. (2011). Role of piRNAs in the Drosophila telomere homeostasis. Mob Genet Elements, 1 (4), 274–278. doi:10.4161/mge.18301

Shpiz, S., Kwon, D., Uneva, A., Kim, M., Klenov, M., Rozovsky, Y., … Kalmykova, A. (2007). Characterization of Drosophila telomeric retroelement TAHRE: transcription, transpositions, and RNAi-based regulation of expression. Mol Biol Evol, 24 (11), 2535–2545. doi:10.1093/molbev/msm205

Shpiz, S., Olovnikov, I., Sergeeva, A., Lavrov, S., Abramov, Y., Savitsky, M., & Kalmykova, A. (2011). Mechanism of the piRNA-mediated silencing of Drosophila telomeric retrotransposons. Nucleic Acids Res, 39 (20), 8703–8711. doi:10.1093/nar/gkr552

Simkin, A., Wong, A., Poh, Y. P., Theurkauf, W. E., & Jensen, J. D. (2013). Recurrent and recent selective sweeps in the piRNA pathway. Evolution, 67 (4), 1081–1090. doi:10.1111/evo.12011

Song, J., Liu, J., Schnakenberg, S. L., Ha, H., Xing, J., & Chen, K. C. (2014). Variation in piRNA and transposable element content in strains of Drosophila melanogaster. Genome Biol Evol, 6 (10), 2786–2798. doi:10.1093/gbe/evu217

Stamatakis, A. (2014). RAxML version 8: a tool for phylogenetic analysis and post-analysis of large phylogenies. Bioinformatics, 30 (9), 1312–1313. doi:10.1093/bioinformatics/btu033

Thomas, J. H., & Schneider, S. (2011). Coevolution of retroelements and tandem zinc finger genes. Genome Res, 21 (11), 1800–1812. doi:10.1101/gr.121749.111

Thurmond, J., Goodman, J. L., Strelets, V. B., Attrill, H., Gramates, L. S., Marygold, S. J., … FlyBase, C. (2019). FlyBase 2.0: the next generation. Nucleic Acids Res, 47(D1), D759–D765. doi:10.1093/nar/gky1003

Traverse, K. L., & Pardue, M. L. (1988). A spontaneously opened ring chromosome of Drosophila melanogaster has acquired He-T DNA sequences at both new telomeres. Proc Natl Acad Sci U S A, 85 (21), 8116–8120. doi:10.1073/pnas.85.21.8116

Villasante, A., Abad, J. P., Planello, R., Mendez-Lago, M., Celniker, S. E., & de Pablos, B. (2007). Drosophila telomeric retrotransposons derived from an ancestral element that was recruited to replace telomerase. Genome Res, 17 (12), 1909–1918. doi:10.1101/gr.6365107

Volff, J. N. (2006). Turning junk into gold: domestication of transposable elements and the creation of new genes in eukaryotes. Bioessays, 28 (9), 913–922. doi:10.1002/bies.20452

Walter, M. F., Biessmann, M. R., Benitez, C., Torok, T., Mason, J. M., & Biessmann, H. (2007). Effects of telomere length in Drosophila melanogaster on life span, fecundity, and fertility. Chromosoma, 116 (1), 41–51. doi:10.1007/s00412-006-0081-5

Wang, L., Barbash, D. A., & Kelleher, E. S. (2019). Divergence of piRNA pathway proteins affects piRNA biogenesis and off-target effects, but not TE transcripts, revealing a hidden robustness to piRNA silencing. bioRxiv.

Wang, L., Dou, K., Moon, S., Tan, F. J., & Zhang, Z. Z. (2018). Hijacking Oogenesis Enables Massive Propagation of LINE and Retroviral Transposons. Cell, 174 (5), 1082–1094 e1012. doi:10.1016/j.cell.2018.06.040

Wang, W., Yoshikawa, M., Han, B. W., Izumi, N., Tomari, Y., Weng, Z., & Zamore, P. D. (2014). The initial uridine of primary piRNAs does not create the tenth adenine that Is the hallmark of secondary piRNAs. Mol Cell, 56 (5), 708–716. doi:10.1016/j.molcel.2014.10.016

Waterhouse, A. M., Procter, J. B., Martin, D. M., Clamp, M., & Barton, G. J. (2009). Jalview Version 2--a multiple sequence alignment editor and analysis workbench. Bioinformatics, 25 (9), 1189–1191. doi:10.1093/bioinformatics/btp033

Wei, K. H., Reddy, H. M., Rathnam, C., Lee, J., Lin, D., Ji, S., … Barbash, D. A. (2017). A Pooled Sequencing Approach Identifies a Candidate Meiotic Driver in Drosophila. Genetics, 206 (1), 451–465. doi:10.1534/genetics.116.197335

Wintersinger, J. A., & Wasmuth, J. D. (2015). Kablammo: an interactive, web-based BLAST results visualizer. Bioinformatics, 31 (8), 1305–1306. doi:10.1093/bioinformatics/btu808

Zhao, K., Cheng, S., Miao, N., Xu, P., Lu, X., Zhang, Y., … Yu, Y. (2019). A Pandas complex adapted for piRNA-guided transposon silencing. bioRxiv.

